# Diffusiophoresis promotes phase separation and transport of biomolecular condensates

**DOI:** 10.1101/2023.07.03.547532

**Authors:** Viet Sang Doan, Ibraheem Alshareedah, Anurag Singh, Priya R. Banerjee, Sangwoo Shin

**Affiliations:** Department of Mechanical and Aerospace Engineering, University at Buffalo, The State University of New York, Buffalo, NY 14260; Department of Physics, University at Buffalo, The State University of New York, Buffalo, NY 14260

## Abstract

The internal microenvironment of a living cell is heterogeneous and comprises a multitude of organelles with distinct biochemistry. Amongst them are biomolecular condensates, which are membrane-less, phase-separated compartments enriched in system-specific proteins and nucleic acids. The heterogeneity of the cell engenders the presence of multiple spatiotemporal gradients in chemistry, charge, concentration, temperature, and pressure. Such thermodynamic gradients can lead to non-equilibrium driving forces for the formation and transport of biomolecular condensates. Here, we report how ion gradients impact the transport processes of biomolecular condensates on the mesoscale and biomolecules on the microscale. Utilizing a microfluidic platform, we demonstrate that the presence of ion concentration gradients can accelerate the transport of biomolecules, including nucleic acids and proteins, via diffusiophoresis. This hydrodynamic transport process allows localized enrichment of biomolecules, thereby promoting the location-specific formation of biomolecular condensates via phase separation. The ion gradients further impart active motility of condensates, allowing them to exhibit enhanced diffusion along the gradient. Coupled with a reentrant phase behavior, the gradient-induced active motility leads to a dynamical redistribution of condensates that ultimately extends their lifetime. Together, our results demonstrate diffusiophoresis as a non-equilibrium thermodynamic force that governs the formation and transport of biomolecular condensates.

---

Biomolecular condensates are a group of non-membraneous organelles that carry out a myriad of intracellular functions including stress response,^1^ intracellular signaling,^2^ and genome organization.^3^ These condensates concentrate organelle-specific proteins and nucleic acids in a spatiotemporal manner to achieve their desired functions in the cell. Segregative transitions, such as phase separation, have been heavily cited as the most plausible mechanism for the formation of biomolecular condensates.^4^ Phase separation is a density transition that occurs through a hierarchy of attractive chain-chain and repulsive chain-solvent interactions leading to the formation of compositionally distinct macromolecular phases.^5^

The spatial location and active transport of biomolecular condensates within the cell are often tightly regulated. For example, P granules, which are RNA- and protein-rich condensates, form near the posterior of the cytoplasm during *C. elegans* zygote polarization.^6^ The asymmetric spatial patterning of P granules has been attributed to the intracellular concentration gradients of proteins and RNA.^7^ Similar observations were made for bacterial PopZ condensates that regulate cell division in bacteria.^8^ PopZ condensates locally form at the poles of bacterial cells. Although not fully understood, it has been hypothesized that PopZ condensates localize to the poles of the bacterial cells due to less crowded chromatin at the poles.^9^ Plant cells provide further examples of the spatially controlled formation of condensates.^10^ These examples and many others suggest that the site-specific localization of condensates within the cell is a prerequisite to their function.^11^ For these reasons, a deeper understanding of the biophysical mechanisms that dictate condensate spatial localization and transport is a topic of significant importance.

The biological cell is intrinsically heterogeneous in space and time, exhibiting several types of thermodynamic gradients such as temperature, pressure, ions, and biological macromolecules.^12,13^ For instance, thermometry studies have shown that the nucleus and other intracellular organelles can lead to local variations in intracellular temperatures due to their behaviors as local heat sources.^14^ Pressure gradients are also evident within the living cells where the variations in the ion concentration lead to both osmotic and hydraulic pressures across the cell.^15^ The presence of gradients often leads to nonequilibrium processes, such as the asymmetric diffusion of molecules and particles up and down the gradients.^16^ Thermophoretic particles migrate to regions of higher (thermophilic) or lower (thermophobic) temperature when a temperature gradient is established.^17^ Charged particles exhibit active transport in the presence of electrical potential gradients, a phenomenon that is known as electrophoresis and used in many technological applications.^18,19^

Likewise, the migration of particles induced by solute gradients is referred to as diffusiophoresis.^20,21^ In a metabolically active cell, it is speculated that diffusiophoresis may facilitate the transport of macromolecular assemblies across the cell cytoplasm.^22^ However, it is unknown whether thermodynamic gradients within the cell can cause nonequilibrium forces to drive the motility of biomolecular condensates in space and time through diffusiophoresis. Importantly, the diffusiophoretic response of biomolecular condensate to a gradient would depend on the interfacial properties of the condensate and the type of gradients present. Therefore, understanding how biomolecular condensates behave in the presence of gradients is critical for elucidating their spatial patterning and active motion within the cell.

In this work, we study the effect of salt concentration gradients on the formation and transport of biomolecular condensates formed by associative phase separation of multivalent disordered proteins and nucleic acids. We postulate two types of effects that a gradient may impart on a biomolecular condensate system: (a) the gradient dictates the regions where the formation of biomolecular condensates via phase separation is favorable, and (b) the gradient leads to active motility of a biomolecular condensate by biasing the condensate diffusion towards a certain direction with respect to the gradient axis. Both of these effects are plausible and may occur concurrently.

To understand these two effects and their interplay with the biophysical properties of condensates, we employ an *in vitro* model system comprised of Arg/Gly-rich multivalent peptide [RGRGG]_5_ (25 amino acids) and a single-stranded homopolymeric DNA [dT]_40_ (40 nucleotides). Arg/Gly-rich disordered protein domains have been shown to drive ribonucleoprotein phase separation with RNA^23^ and are present in a large percentage of the RNA-binding and condensate-forming proteome.^24^ These positively charged multivalent domains undergo phase separation with nucleic acids through a combination of their attractive electrostatic forces with the negatively charged backbone of nucleic acids and cation-π and π-π interactions with nucleobases.^25–27^ Several recent studies have characterized the phase behavior and material properties of RGG domains with ssDNA and RNA.^23,27–30^ Importantly, RGG-nucleic acid (NA) phase separation is reentrant, meaning that phase separation is only favored within a finite window of mixing ratios that are usually centered around the stoichiometric charge-balanced mixture composition.^31^ The stoichiometry of RGG-NA mixtures also dictates the interfacial charge of these condensates via a charge inversion mechanism.^25,32,33^ The tunable phase behavior and interfacial properties of RGG-NA condensates make them suitable systems for studying the effect of ion gradients on condensate motility.

Using a controlled microfluidic setup alongside fluorescence microscopy, we find that concentration gradients of the peptide and the ssDNA promote spatially patterned condensate formation in specific regions along the salt gradient. We show that the formation of condensates is more robust in the presence of salt gradients due to the local enrichment of the oppositely charged biomolecules via diffusiophoresis. After their formation, the presence of the salt gradient enhances the motility of condensates along the gradient and establishes their active transport that is dependent on the condensate surface charge. Microfluidics studies on peptide and ssDNA solutions reveal that spatial patterning of condensates is dictated by the location-specific concentration and stoichiometry of peptide and ssDNA mixtures. These results are extended to other biomolecular systems where phase separation is driven by obligate heterotypic electrostatic interactions, such as in the mixtures of cationic nucleic acid binding protein protamine and homopolymeric RNA, poly(rU). Overall, our results show that the controlled spatial localization of biomolecular condensates can be spontaneously achieved with macromolecular and ionic concentration gradients. Furthermore, the active motility of condensates in the presence of salt gradients can add additional control over their localization. Together, these findings shed light on the role of ion and chemical gradients in controlling the localization and transport of biomolecular condensates and highlight diffusiophoresis as a plausible mechanism of localization control of biomolecular condensates within cells.

## Phase separation is promoted in the presence of salt gradients

To investigate the associative phase separation of ssDNAs and cationic polypeptides in the presence of salt gradients, we employ a controlled microfluidic platform (**Fig. 1**). In an H-shaped microfluidic channel made out of a UV-curable epoxy (NOA-81), we initially fill the entire channel with the polypeptide [RGRGG]_5_ (0.5 mg/ml; 0.17 mM) in Tris buffer (pH 7.5) with additional NaCl of concentration *c*_1_ (**Fig. 1a** and **Supplementary Fig. S1**). Then, along the left reservoir channel, we supply an oppositely charged ssDNA [dT]_40_ (0.63 mg/ml; 0.13 mM) in Tris buffer (pH 7.5) with NaCl of concentration *c*_2_. Finally, the [RGRGG]_5_ in Tris buffer with NaCl of concentration of *c*_1_ is injected into the right reservoir. A thin hydrogel membrane (polyethylene glycol diacrylate, PEGDA) is patterned at the right end of the horizontal channel to suppress undesired advective flows in the horizontal channel while allowing the diffusive transport of the molecular solutes.^34,35^ This ensures that only salt gradients are established while keeping the buffer conditions identical throughout the channel. The PEGDA membrane also suppresses [dT]_40_ and [RGRGG]_5_ from permeating through the gel. The side channels thus act as reservoirs that provide and sustain gradients of salts and biomolecules within the center channel. Therefore, the experiments are performed such that a finite amount of initially present polypeptide is gradually diffused out while ssDNA is continuously provided through the left reservoir in the presence (or absence) of salt gradients. Overall, our microfluidic setup can robustly create thermodynamic gradients of ions and biological macromolecules mimicking heterogeneous subcellular microenvironment.^36^

**Figure 1.**
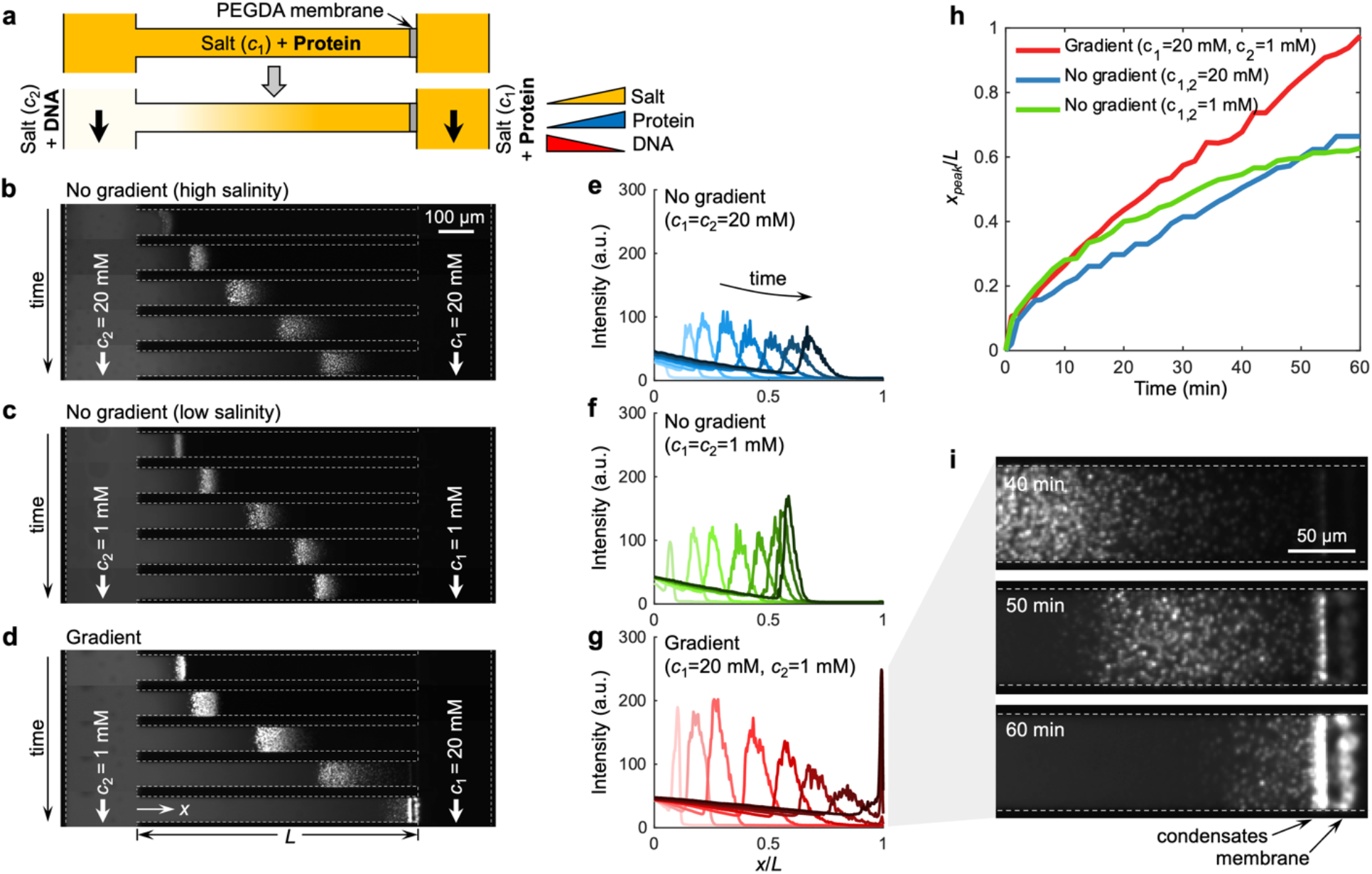
Salt gradients promote the transport and phase separation of biomolecular condensates. (**a**) The experimental setup employs an H-shaped microfluidic device with a PEGDA hydrogel membrane patterned at the right end of the center horizontal channel (length *L* = 800 µm) that allows the solute species to diffuse into the channel but prevents fluid flow. The entire channel is first flushed with polypeptide ([RGRGG]5, 0.5 mg/ml) suspended in Tris buffer (pH 7.5) and NaCl of concentration *c*_1_. After that, ssDNAs ([dT]40, 0.63 mg/ml) in Tris buffer (pH 7.5) with NaCl of concentration *c*_2_ is infused along the left reservoir channel, whereas the same polypeptide in Tris buffer (pH 7.5) with NaCl of concentration of *c*_1_ is injected into the right reservoir. (**b-d**) Fluorescence image sequences and (**e-g**) their corresponding width-averaged intensity distribution of the [dT]40-[RGRGG]5 condensates over a period of 60 min. The NaCl concentrations are (**b**,**e**) *c*_1_ = *c*_2_ = 20 mM, (**c**,**f**) *c*_1_ = *c*_2_ = 1 mM, and (**d**,**g**) *c*_1_ = 20 mM, *c*_2_ = 1 mM. The timestamps for the images in (**b-d**) are 1, 5, 20, 40, and 60 min, and for the plots in (**e-g**) are 1, 5, 10, 20, 30, 40, 50, and 60 min. (**h**) The location of the peak intensity *x*_*peak*_ over time for different NaCl concentrations. (**i**) Close-up image series of the condensates near the PEGDA membrane from (**d**,**g**), showing localized accumulation.

We monitor the phase separation and transport of [dT]_40_ and [RGRGG]_5_ using fluorescence microscopy, as shown by the time-lapse images in **Figs. 1b-d** (**Supplementary Movie S1**) and their corresponding fluorescence intensity profiles in **Figs. 1e-g**. The condensates are visualized using Cy5-labeled [dT]_40_ (1.2 mol% of the total [dT]_40_). Once [dT]_40_ is introduced to the left reservoir, it starts to interact with [RGRGG]_5_ and undergoes phase separation via multivalent heterotypic interactions.^30^ These condensates are polydispersed submicron-sized coacervates with mean diameters ranging between 230–660 nm depending on the buffer salt concentration (**Supplementary Fig. S2**).

Similar to complex coacervates, RGG-NA phase separation is suppressed with an increasing salt concentration in the buffer since electrostatic interactions are one of the major drivers of phase separation of the mixture.^25,33^ Consistent with this, we observe that with less NaCl in the background (*c*_1_ = *c*_2_ = 1 mM; **Figs. 1c,f**), phase separation occurs stronger compared to the higher salinity case (*c*_1_ = *c*_2_ = 20 mM; **Figs. 1b,e**) as evidenced by the higher fluorescence intensity. What is remarkable is that when a salinity gradient is introduced (*c*_1_ = 20 mM and *c*_2_ = 1 mM; **Figs. 1d,g**), the fluorescence intensity becomes even stronger, despite the overall salinity being higher than the low salinity case (*c*_1_ = *c*_2_ = 1 mM) and the similarity in biomolecular concentrations. With NaCl gradients, the overall intensities are significantly higher than the experiments performed without the gradients (**Fig. 1g**). These results indicate that NaCl gradients promote peptide-ssDNA coacervation.

We further observe that peptide-ssDNA condensates are not only formed locally inside the channel but also display wave-like patterns that move along the channel. Notably, the wave speed is faster and moves further down the channel when the NaCl gradient is present, as shown in **Fig. 1h**, where we track the position of the wave peaks *x*_*peak*_ over time. We note that the condensates under NaCl gradients eventually reach the far right end of the channel where the PEGDA membrane is present. The condensates subsequently accumulate adjacent to the membrane, displaying an extremely localized distribution (**Fig. 1i**).

We posit that the observed directional migration of the condensates in the NaCl gradients is caused by diffusiophoresis, which describes a spontaneous migration of charged colloidal particles driven by ionic solute gradients.^37^ Given that the concentration of [dT]_40_ in the left reservoir is higher than the initial [RGRGG]_5_ concentration in the center channel, the condensates are generally rich in [dT]_40_. These condensates are expected to be negatively charged with excess [dT]_40_ populated preferentially at the surface of the condensate.^38^ Indeed, the electrophoretic light scattering measurements of the condensates at the mixture ratio of [dT]_40_:[RGRGG]_5_ = 1.25:1 confirms the negative surface charge with a zeta potential *ζ* = –27.4 mV (**Supplementary Fig. S3**). These negatively charged condensates migrating toward the higher NaCl concentration side are consistent with previously reported diffusiophoresis of negatively charged colloids in NaCl gradients.^39–41^ The logarithmic sensing of diffusiophoresis, i.e., the diffusiophoretic velocity dependent on the gradient of the logarithm of the salt concentration (**Supplementary Fig. S4**), further strengthens the nature of the directional migration being diffusiophoresis.^42,43^

We next employed image analysis to track individual condensates within the wave in order to understand the detailed dynamics of the condensate wave motion in **Figs. 2a-c** (**Supplementary Movie S2**). We plot a kymograph showing the spatiotemporal dynamics of the wave and the condensates. In the absence of NaCl gradient, we find that individual condensate trajectories are stochastic and do not exhibit a directional bias, appearing as flat tracks as a function of time (**Figs. 2a,b**). This indicates that the condensate wave is propagating by the dissolution of condensates at the rear side of the wave and the formation of new condensates at the wavefront. Contrastingly, in the presence of salt gradients, the advective trajectories of multiple condensates show that they actively migrate toward the higher NaCl concentration side (**Fig. 2c**). This active migration of trailing condensates effectively delays their dissolution. Overall, these findings indicate that NaCl gradients not only enhance condensate formation compared to the uniform salt conditions, as indicated by the intensity signal being stronger in **Fig. 2c**, but also lead to a directional migration of peptide-ssDNA condensates toward higher NaCl concentrations.

**Figure 2.**
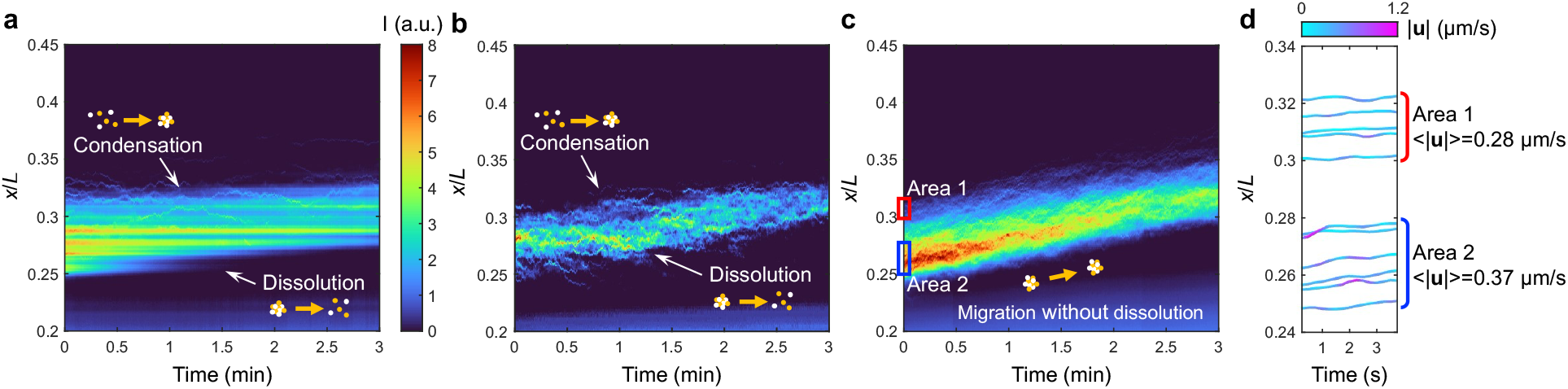
Spatiotemporal dynamics of biomolecular condensates reveal the influences of diffusiophoresis on their formation, transport, and dissolution. (**a**,**b**) Kymographs of the fluorescence signal of the [dT]40-[RGRGG]5 condensates without the presence of NaCl gradient [**(a)** *c*_1_ = *c*_2_ = 20 mM; **(b)** *c*_1_ = *c*_2_ = 1 mM]. The waves of the condensate are formed due to simultaneous dissolution near the front and condensation near the rear. (**c**) With NaCl gradients (*c*_1_ = 20 mM, *c*_2_ = 1 mM), the formation of the droplets with the migration toward the higher salt concentration prevents the condensates from dissolution. The color code represents the fluorescence intensity *I*. (**d**) The trajectories of the individual condensates found near the front (*x*/*L* ~ 0.31, area 1) and rear (*x*/*L* ~ 0.26, area 2) of the wave in (**c**). The color code represents the velocity magnitude |***u***|. 〈|***u***|〉 represents the mean of the time-averaged velocity magnitude of multiple condensates in each region.

## The promotion of phase separation is a result of the local enrichment of biomolecules driven by diffusiophoresis

Diffusiophoresis alters the transport of not only charged colloidal particles but also charged molecular species.^44,45^ Given that [dT]_40_ and [RGRGG]_5_ are oppositely charged, we speculate that the stronger phase separation of the [dT]_40_-[RGRGG]_5_ mixtures under NaCl gradients is due to the local enrichment of the individual biomolecules driven by differential diffusiophoretic transport. To test this thesis, we conducted similar microfluidic experiments with only either [dT]_40_ or [RGRGG]_5_ present in the same setup with or without the NaCl gradients (**Figs. 3a-d**). In these experiments, not only [dT]_40_ is fluorescently tagged (1.2 mol% of the total [dT]_40_), but also [RGRGG]_5_ is tagged with Alexa-488 fluorophores (3.7 mol% of the total [RGRGG]_5_). Both The migration of [dT]_40_ entering from the left reservoir without the NaCl gradient is driven solely by diffusion, which shows a monotonically decaying distribution along the channel (**Fig. 3a**). In contrast, with the presence of the NaCl gradient, negatively charged [dT]_40_ molecules experience a non-monotonic distribution where the molecules locally accumulate as they diffuse down the channel (**Fig. 3b**).

**Figure 3.**
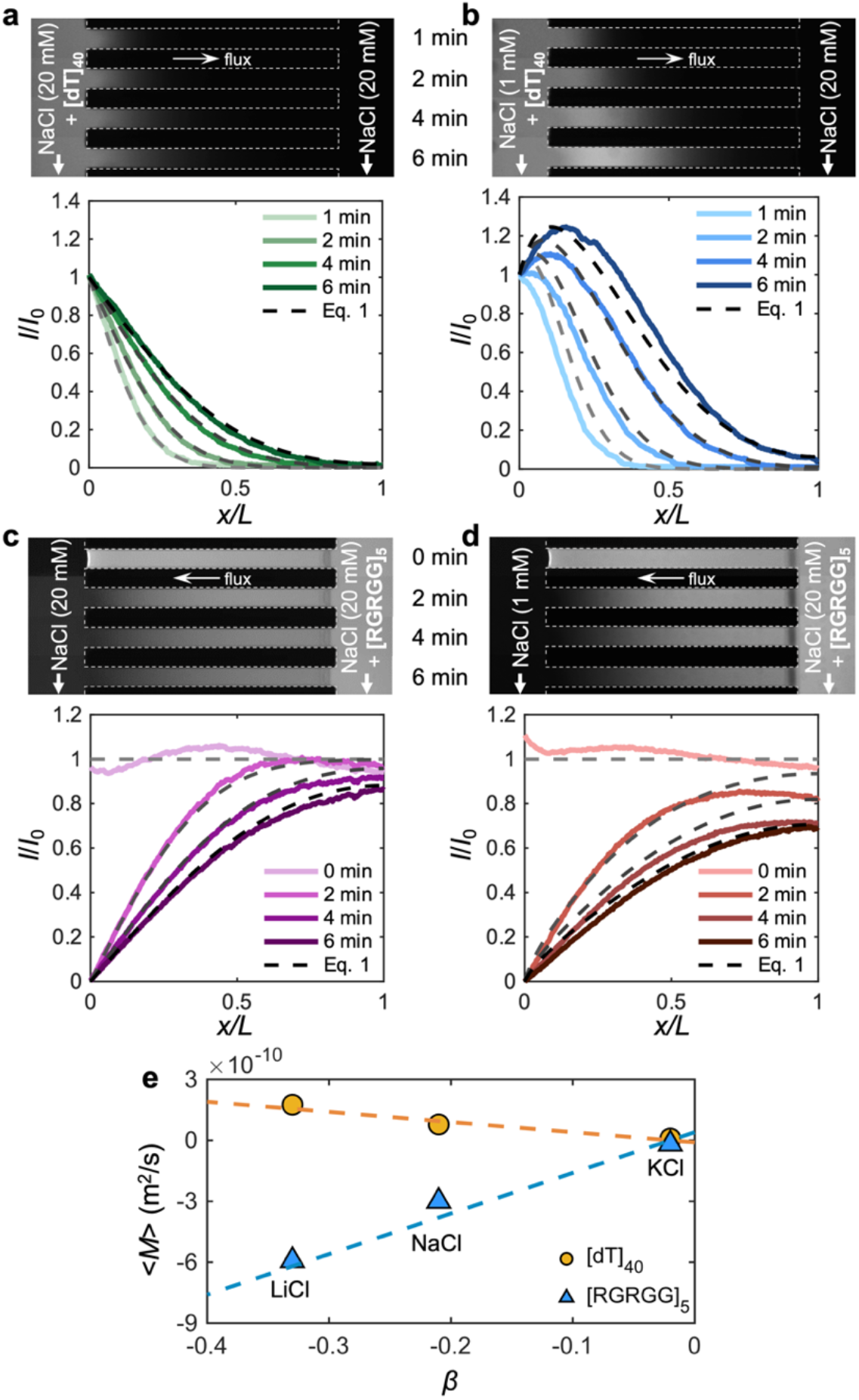
Diffusiophoresis enhances the transport of biomolecules. (**a-d**) Diffusion experiments for individual biomolecules where either only (**a**,**b**) [dT]40 or (**c**,**d**) [RGRGG]5 are introduced to the side channel in the (**a**,**c**) absence (*c*_1_ = *c*_2_ = 20 mM) or (**b**,**d**) presence or NaCl gradients (*c*_1_ = 20 mM, *c*_2_ = 1 mM). For these experiments, we used Cy5-labeled [dT]40 (1.2 mol% of the total [dT]40) and Alexa488-labeled [RGRGG]5 (3.7 mol% of the total [RGRGG]5). The top panels are the fluorescence images and the bottom panels are width-averaged intensity profiles. The dashed curves are generated using equation 1. (**e**) The time-averaged diffusiophoretic mobility <*M*> for [dT]40 (yellow circles) and [RGRGG]5 (blue triangles) molecules in monovalent chloride salts of diffusivity contrast *β*. The dash lines represent the best linear fitting of the mobility and *β*. The mobility tends to zero as *β* approaches zero, indicating a negligible effect of chemiphoresis.

This behavior is likely due to the logarithmic nature of diffusiophoresis.^42^ As the diffusiophoretic velocity (*u*_*d*_) scales with the gradient of the logarithm of the salt concentration (*c*), viz., *u*_*d*_ ~ *∂*_*x*_ ln *c*, the velocity is influenced by the absolute salt concentration where the biomolecules undergoing diffusiophoresis slow down as they migrate toward higher salt concentrations, leading to their accumulation. This peculiar feature has been observed in a variety of biocolloids including bacterial cells,^46^ liposomes,^47,48^, DNAs,^49,50^ and proteins.^44^ Notably, Riback et al. recently reported similar wave-like advective migration of ribosomal RNAs, a nuclear biomolecular condensate associated with ribosome biogenesis,^51^ for which the migration is speculated to be due to the viscoelasticity gradients present in the nucleolus.^52^ Finally, similar to the ssDNA migration, positively charged [RGRGG]_5_ molecules also experience enhanced transport via diffusiophoresis, but they migrate down the gradient due to the reversed charge polarity (**Figs. 3c,d**).

To quantify the diffusion coefficient and the diffusiophoretic mobility of [dT]_40_ and [RGRGG]_5_ moving up and down the NaCl gradient from the fluorescence intensity distributions, we consider a one-dimensional transport equation for the individual biomolecules,^39^ which reads

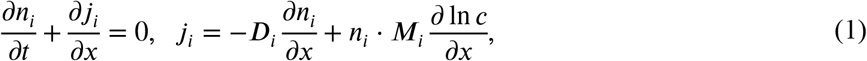

where *n*_*i*_ is the mass concentration, *j*_*i*_ is the mass flux, *D*_*i*_ is the diffusivity, and *M*_*i*_ is the diffusiophoretic mobility of the biomolecular species *i* (*i* ∈ {DNA (*D*), polypeptide (*P*)}). The mobility *M*_*i*_ quantifies the tendency of a biomolecule’s migration in response to local concentration fields *c*. The second term on the right-hand side for the flux equation is the phoretic advection driven by the salt concentration gradient. Here, [dT]_40_ molecules diffuse into the channel with the bulk diffusivity of *D*_*D*_ = 1.2×10^−10^ m^2^/s (**Fig. 3b**). With diffusiophoresis, an advective velocity *u*_*d*_ = *M*_*i*_*∂*_*x*_ ln *c* is added to the transport of [dT]_40_ molecules, boosting the migration with additional mobility of *M*_*R*_ = 0.78×10^−10^ m^2^/s (**Fig. 3a**). Given the units for mobility being identical to the diffusivity, diffusiophoretic mobility can alternatively be viewed effectively as an enhanced diffusivity.^53,54^

For [RGRGG]_5_, which diffuses out through the left reservoir, the diffusivity without NaCl gradient is estimated as *D*_*P*_ = 2.3×10^−10^ m^2^/s (**Fig. 3c**). As the NaCl gradient is introduced, [RGRGG]_5_ diffuses out of the channel much faster, accelerated by the diffusiophoretic mobility of *M*_*P*_ = –0.3×10^−10^ m^2^/s (**Fig. 3d**). The negative sign indicates that [RGRGG]_5_ migrates down the NaCl gradient. Unlike [dT]_40_ migrating up the concentration gradients by which the molecules locally accumulate, the logarithmic dependence now causes dispersion of [RGRGG]_5_ toward the lower salt concentration front. Therefore, while [dT]_40_ is enriched from lower to higher NaCl concentration, [RGRGG]_5_ is quickly brought from higher to lower NaCl concentration. The opposing actions of diffusiophoresis on [dT]_40_ and [RGRGG]_5_ effectively create an attractive interaction over a longer distance, where the length scale is effectively set by the salt diffusion length.^55^ This local enrichment of both of these oppositely charged biomolecules directly promotes the phase separation of [dT]_40_-[RGRGG]_5_ condensates.

We further confirm the diffusiophoretic transport of [dT]_40_ and [RGRGG]_5_ by performing experiments in other similar monovalent chloride salts, namely LiCl and KCl. For charged molecular species, their diffusiophoretic mobility is linearly proportional to the diffusivity contrast factor, *β* = (*D*_+_ – *D*_–_)/(*D*_+_ + *D*_–_), where *D*_+_ and *D*_–_ are the diffusivity of cations and anions, respectively.^56^ *β* sets the magnitude of the diffusion potential that provides the electrokinetic driving force for diffusiophoresis. Indeed, we observe a linear dependence of the mobility on *β* for both [dT]_40_ and [RGRGG]_5_ (**Fig. 3e** and **Supplementary Fig. S5**). Also, their distinct mobility signs confirm once again that the transport of oppositely charged biomolecules is accelerated toward each other by diffusiophoresis, providing a non-equilibrium driving force to phase separate. These results collectively highlight the importance of non-equilibrium interactions of ionic species with biomolecules and their condensates. The equilibrium electrostatics of the monovalent cations Na^+^, K^+^, and Li^+^ are more or less identical, yet their non-equilibrium electrokinetics vary significantly, as manifested by *β*.

## Formation and propagation of the condensate waves are the results of reentrant phase behavior and non-linear diffusiophoresis

Regardless of the presence of NaCl gradients, the biomolecular condensates are distributed in a wave-like fashion where the condensates are formed only within a narrow region in the channel (although the wave speed is significantly influenced by the NaCl gradients), as shown in the intensity profiles in both cases (**Figs. 1e-g**). These profiles are replotted in **Figs. 4a,b** where we now delineate the phase-separated regions (solid curves) from the regions of no visible condensates (dashed curves). This behavior in which the phase separation occurs locally is reminiscent of the reentrant phase behavior where the phase separation takes place only within a specific range of biomolecule stoichiometry.^31^ Since [dT]_40_ enters from the left while [RGRGG]_5_ migrates from the right, the concentration ratio of the two biomolecules (i.e., *n*_*D*_/*n*_*P*_ where *n*_*D*_ and *n*_*P*_ are the mass concentration of [dT]_40_ and [RGRGG]_5_, respectively) gradually decreases along the channel. Such a distribution results in the localized zone of phase separation that is set by a finite window of *n*_*D*_/*n*_*P*_.

**Figure 4.**
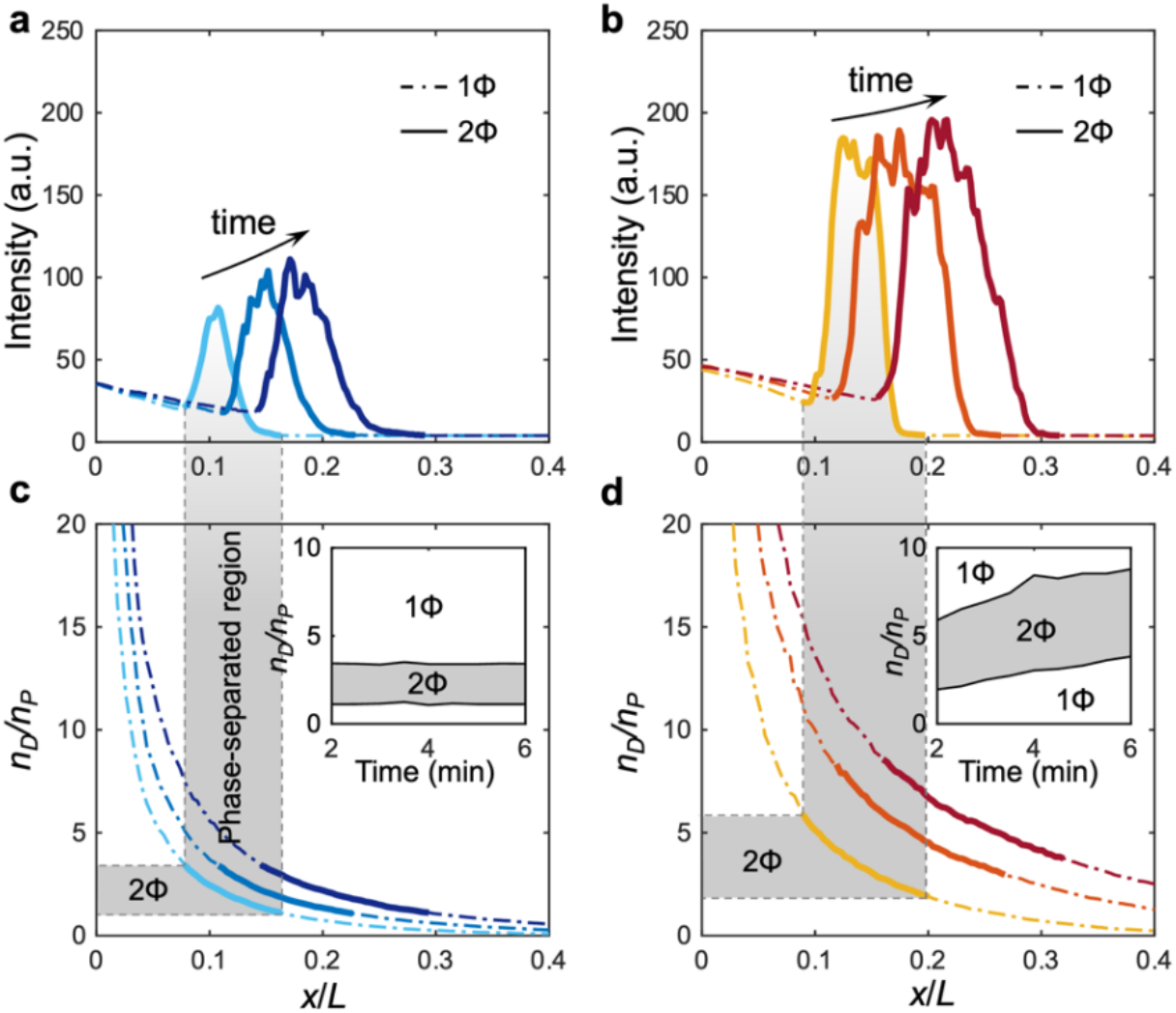
Reentrant phase behavior results in wave-like distribution of condensates that move along the gradients. (**a**,**b**) Fluorescence intensity profiles taken from the experiments in **Figs. 1b** (*c*_1_ = *c*_2_ = 20 mM) and **1d** (*c*_1_ = 20 mM, *c*_2_ = 1 mM). The phase-separated regions are depicted by the solid curves whereas the homogeneous regions are indicated by the dash-dot curves. The timestamps are 2, 4, and 6 min. (**c**,**d**) The concentration ratio of [dT]40 and [RGRGG]5 (*nD*/*nP*) reconstructed from the individual biomolecule experiments in **Figs. 3a-d**. Insets represent the phase boundaries identified from the intensity plots. As an example, the phase-separated region at 2 min is indicated by the translucent gray bar for a guide to the eye.

From the intensity distributions for the individual biomolecules presented in **Figs. 3a-d**, by taking the ratio of the two, we can plot the mixture composition ratio *n*_*D*_/*n*_*P*_ versus *x*/*L* (**Figs. 4c,d**). The concentrations of the biomolecules, *n*_*D*_ and *n*_*P*_, are assumed to be linearly proportional to the fluorescence intensity, which is a valid assumption in dilute systems.^57^ As we compare the reconstructed *n*_*D*_/*n*_*P*_ plot (**Fig. 4c**) with the condensate distribution (**Figs. 4a,b**), we indeed observe that the range of *n*_*D*_/*n*_*P*_ over which the phase separation occurs remains more or less constant over time when the NaCl concentration is uniform throughout the channel. This is also shown in the inset of **Fig. 4c**, where we explicitly plot the phase separation range of *n*_*D*_/*n*_*P*_ (gray region). As [dT]_40_ and [RGRGG]_5_ diffuse toward each other, the local concentrations of the two biomolecules, and thus the ratio of the two (*n*_*D*_/*n*_*P*_), change dynamically in space and time. This gradually shifts the phase-separated region in the positive *x*-direction by forming new condensates in the leading front and dissolving the existing condensates in the trailing end (**Fig. 5a**), thus creating a wave-like condensate profile. This is also captured in the kymograph (**Fig. 2a**), where the condensate streaks that fade in and out indicate, respectively, condensation and dissolution. Despite the dynamically changing *n*_*D*_/*n*_*P*_, the condensates are always formed within a fixed range of stoichiometry (*n*_*D*_/*n*_*P*_), as also indicated by the constant width of the gray region in the inset of **Fig. 4c**. This suggests that the formation and dissolution of the condensates occur much faster than the migration of the phase-separation region, thus indicating that the phase separation occurs in a quasi-equilibrium manner. Specifically, our microfluidics experiments indicate that condensates form when 1.0 < *n*_*D*_/*n*_*P*_ < 3.3.

**Figure 5.**
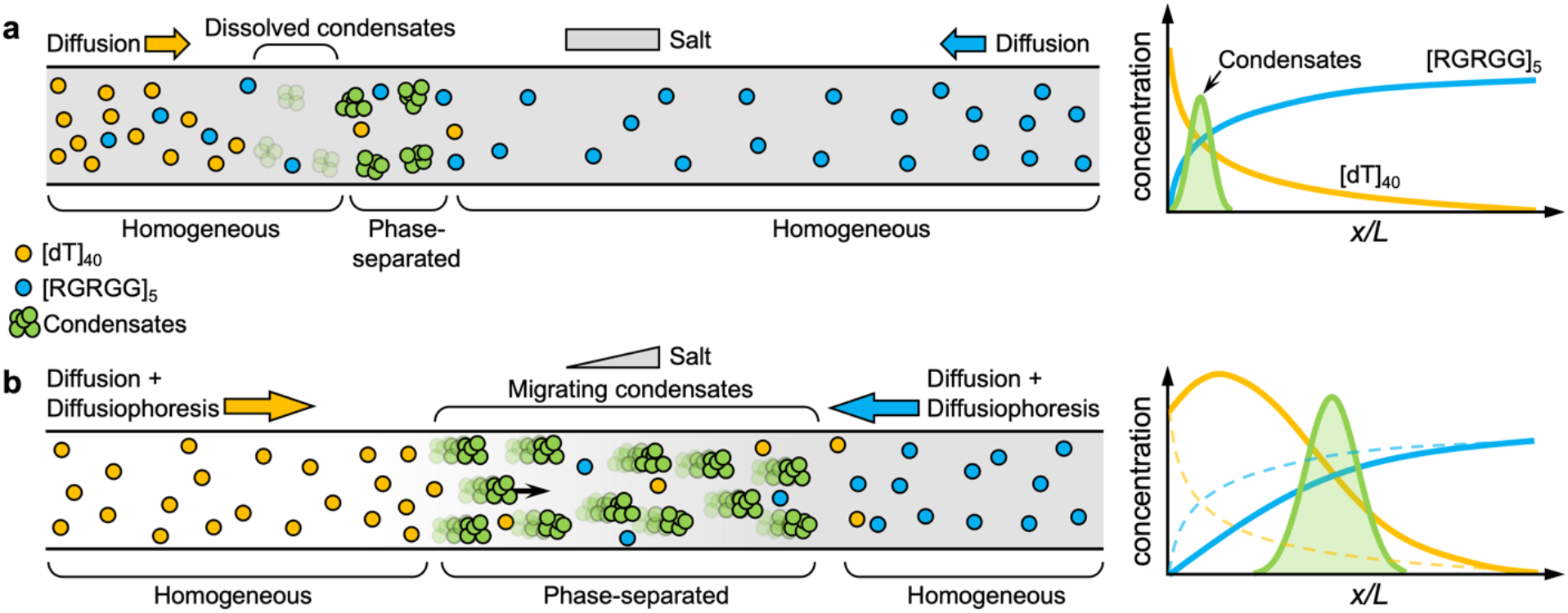
Enhanced phase separation and transport of condensates through diffusiophoresis. (**a**) A schematic diagram showing how reentrant phase separation underlies the wave-like patterning of peptide-ssDNA condensates. In the absence of a salt gradient, both [dT]40 and [RGRGG]5 diffuse from both sides, leading to the formation of biomolecular condensates. As a result of the reentrant phase behavior, these condensates experience dissolution and condensation, forming wave-like profiles as depicted in the plot on the right-hand side (see the experimental data in **Fig. 1b**). (**b**) In the presence of a salt gradient, the transport of biomolecules is enhanced by diffusiophoresis, which increases their local concentration and thus promotes their phase separation. In addition, salt gradients further impart directional motility to the phase-separated condensates, thereby extending their lifetime.

To confirm the quasi-equilibrium nature of the condensate formation, we performed turbidity measurements of [dT]_40_-[RGRGG]_5_ mixtures under equilibrium by quantifying the amount of light scattered from the condensates at a given wavelength (350 nm).^25^ Plotting turbidity as a function of mixture composition *n*_*D*_/*n*_*P*_ shows that condensates form between 0.28 < *n*_*D*_/*n*_*P*_ < 3.5 (**Supplementary Fig. S6**). The wider reentrant window may be attributed to the difference in the measurement methods, where the smaller spatial resolution of the turbidity measurements compared to optical microscopy will likely detect smaller clusters that form at stoichiometries far from the optimal conditions.^58^ Nonetheless, the agreement of the two methods suggests that the phase separation behavior under non-equilibrium is dictated by local equilibrium thermodynamics. This is consistent with a recent report that suggested that local thermodynamic equilibrium governs phase separation in living cell cytoplasm despite their inherent non-equilibrium nature.^59^

In the case of NaCl gradients (**Figs. 4b,d**), we observe that the phase-separation range gradually broadens, as also indicated by the widening of the gray region in the inset of **Fig. 4d**. This is attributed to the increase in the local biomolecule concentration driven by diffusiophoresis (i.e., molecular diffusiophoresis), where the local enrichment of biomolecules by diffusiophoresis broadens the two-phase coexistence window. This behavior is consistent with the previous observations under equilibrium conditions with a similar biomolecular system comprised of RNAs and proteins, where the increase in the absolute concentration of RNA [poly(rU)] and cationic proteins results in a wider range of mixture stoichiometry that promote phase separation in equilibrium environments.^25^ In our system, [dT]_40_ and [RGRGG]_5_ undergo diffusiophoresis, causing a local increase in their concentrations in the presence of a NaCl gradient and directly expanding the stoichiometric range of the two-phase region. Unlike the condensates formed without the NaCl gradient, the condensates under the non-equilibrium environments effectively move along with NaCl gradient via diffusiophoresis (i.e., condensate diffusiophoresis). This active, directional motility allows the condensates to keep up their position within the moving front of the phase-separated region (**Fig. 2c**).

The spatial variation within the given stoichiometry window suggests that the condensates’ surface charge may also vary spatially since the surface charge, and thus the diffusiophoretic mobility, of the condensates is governed by the mixture stoichiometry.^38^ To confirm this, we performed electrophoretic mobility measurements of condensates at varying mixture compositions (**Supplementary Fig. S3**), which showed that the zeta potential of the condensates changes from –17.2 mV when *n*_*D*_/*n*_*P*_ = 1 to –32.3 mV when *n*_*D*_/*n*_*P*_ = 3. This zeta potential variation translates to the mobility difference from *M*_*P*_ = 1.0×10^−10^ m^2^/s (*n*_*D*_/*n*_*P*_ = 1) to *M*_*P*_ = 2.4×10^−10^ m^2^/s (*n*_*D*_/*n*_*P*_ = 3). This implies that, apart from the logarithmic dependence, the condensate will experience slower diffusiophoresis near the front of the wave, causing even more accumulation of the condensates. This is evidenced by tracking the position of the individual condensates shown in **Fig. 2c**, where the leading condensates near the front of the wave move considerably slower (~0.28 µm/s) than the trailing condensates at the rear (~0.37 µm/s). Therefore, the simultaneous action of logarithmic sensing and stoichiometry dependence of diffusiophoresis, combined with an enlarged reentrant phase separation window, makes the trailing condensates catch up with the slower leading condensates, which would otherwise have dissolved, reinforcing the local accumulation and migration of the condensates along the salt gradient (**Fig. 5b**).

We note that the condensate migration velocity (~0.1 µm/s) agrees quantitatively with the theoretical estimates for the diffusiophoretic velocity of charged rigid spheres of similar zeta potential.^60,61^ This suggests that these condensates behave close to a solid, where liquid-like phoretic transport, i.e., Marangoni propulsion, is negligible. This is likely due to the viscoelastic nature of [dT]_40_-[RGRGG]_5_ condensates.^30,33,62^ For highly viscous liquid drops, the viscous shear inside the drop should dominate over the viscous shear acting in the exterior, yielding the condition (*λ*/*R*) · (*ηc* /*η*) ≫ 1, where *λ* ~ 1 nm is the Debye layer thickness, *R* ~ 240 nm is the condensate radius, *η* ~ 1 mPa·s is the dynamic viscosity of the external solution, and *ηc* is the dynamic viscosity of the condensates. Based on our previous measurements of similar peptide-nucleic acid condensates,^30,33^ the viscosity *ηc* ranges between 1 − 10 Pa·s, indeed satisfying the above condition. The overall migration of the condensates being identical regardless of their size, shown by the kymographs in **Fig. 2c**, also suggests that their motion is dictated by diffusiophoresis^39^ rather than the Marangoni effect, which strongly depends on the drop size.^63^

Furthermore, while one may reason that the gradients of [dT]_40_ and [RGRGG]_5_ may also induce the diffusiophoresis of the condensates, this is likely to be negligible due to the biomolecules being dilute while the background solutes are relatively concentrated. Given that the hydrodynamic radius of [dT]_40_ and [RGRGG]_5_ is around 1–2 nm, which is comparable to the Debye layer thickness, chemiphoresis should be insignificant. Therefore, we can expect that condensate diffusiophoresis by [dT]_40_ or [RGRGG]_5_ gradients are mainly driven by electrophoresis. For multi-species (electro)diffusiophoresis, one may express the velocity as 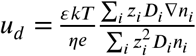, where *ε* is the permittivity, *k*_*B*_ is the Boltzmann constant, *T* is the temperature, *e* is the elementary charge, and *z* is the charge number.^60,61^ Taking the summation over all the known background solute species including the buffer, the velocity is estimated to be less than *ud* = 0.1 nm/s, which is indeed negligible compared to the condensate diffusiophoretic velocity driven by salt gradients.

## Discussion

A number of recent studies have identified that a variety of dynamical processes of biomolecular condensates are associated with thermodynamic activity gradients. For instance, concentration and temperature gradients established by evaporation at liquid-vapor interfaces are shown to promote phase separation in various biomolecular systems.^64,65^ These gradients may further segregate biomolecular condensates by convective transport processes such as Marangoni or gravity-driven flows, having implications in prebiotic compartmentalization and localization.^64,65^ Self-generated chemical gradients via enzymatic reactions can lead to a dynamical response of the condensates. Testa et al. showed that pH gradients established by the enzymatic reactions of urease-containing condensates could induce hydrodynamic flow within and around the condensates via Marangoni flow.^66^ Similar enzymatic-driven gradients may also enable freely suspended condensates to self-propel.^67^ It was also recently predicted that the activity gradients in the nucleus expedite nucleolar coalescence, which helps in the positioning of nucleolus towards the nuclear periphery, a key factor in the localized organization of the nucleus.^3,68^ Other types of gradients, such as protein gradients, have been proposed to drive the localization of biomolecular condensates, such as the asymmetric patterning of P granules in germline polarized cells due to an underlying MEG3 protein gradient.^7^ Chromatin density gradients have also been suggested to drive the local enrichment of PopZ condensates on the poles of bacterial cells.^8^ Therefore, thermodynamic gradients are expected to play a ubiquitous role in regulating the formation and localization of biomolecular condensates in space and time.

In this work, we provided an elaborate study on the effect of salt gradients on biomolecular condensates formed by RGG repeat polypeptides and ssDNA. Using a controlled microfluidic platform, we showed that heterotypic condensates form non-monotonic patterning along salt and biomolecular gradients due to the reentrant nature of their phase separation. We further delineated two effects of salt gradients on RGG-ssDNA condensates that are both driven by diffusiophoresis on different length scales (molecular and mesoscale). The first is that salt gradients enhance the formation of condensates due to their non-linear effect on the diffusiophoresis of the individual protein and ssDNA molecules. The second effect is that salt gradients can impart active migration of the charged condensates, which is controlled by the stoichiometry of peptide-ssDNA mixtures. Both of these effects can have important implications in living cells and can provide multiple levels of localization control over biomolecular condensates by either dictating the spatial location where the condensate formation is favored or by enhancing the diffusion of preformed condensates towards a particular location with respect to the underlying gradient.

More generally, diffusiophoresis, a non-equilibrium driving force generated by (electro)chemical potential gradients, can arise in a wide range of biomolecular condensates. Regardless of whether the biomolecules are negatively or positively charged, natural or synthetic, the individual constituents and the associated condensates can experience diffusiophoresis. We further observe that diffusiophoresis can promote the formation and transport of [RGRGG]_5_-rich, i.e., positively charged, [dT]_40_-[RGRGG]_5_ or poly(rU)-protamine condensates in NaCl gradients (**Supplementary Fig. S7**). Recent studies on diffusiophoresis with a variety of other biomolecules and biocolloids, such as membrane proteins,^69^ exosomes,^48^ bacteria,^46,70^ and blood cells,^71^ or other phase-separating systems^72^ suggest the ubiquity of diffusiophoresis in biological systems where chemical gradients are ever-present, providing suitable conditions for diffusiophoresis to arise.^73^ The presence of a concentrated, active, and nonuniform mixture of proteins, nucleic acids, metabolites, and ions such as ATP, Na^+^, K^+^, and Ca^2+^ inside the cytoplasm makes the movement of particles and molecules subjected to various gradients. For instance, the interdiffusion of equimolar multicomponent solutes of Na^+^ and K^+^ can also induce significant diffusiophoresis and enhanced phase separation (**Supplementary Fig. S8**).

Moreover, the gradient-induced changes in the microenvironment may lead to different dynamical and non-equilibrium properties of the biomolecular condensates. Our work shows that diffusiophoresis is an important phenomenon for the formation and regulation of biomolecular condensates. The methods we presented here can be utilized to study the effect of diffusiophoresis on many levels beyond salt gradients, including other types of biochemical gradients such as ligands, protein/nucleic acid, crowding, enzymes, and other types of co-solutes. Such studies can illuminate our understanding of the spatiotemporal regulation of biomolecular condensates within the active microenvironment of a living cell and can enable technology development toward synthetic membrane-less organelles with precise localization and functionalities.

## Acknowledgments

This work was supported by grants from the National Science Foundation (2223737 and 2237177 to S.S.) and the National Institute of General Medical Sciences of the National Institutes of Health (R35 GM138186 to P.R.B.). The authors acknowledge members of the Banerjee and Shin labs for valuable discussions during different stages of the manuscript preparation.

## Author Contributions

Conceptualization: P.R.B. and S.S.; Methodology: V.S.D., I.A., P.R.B., and S.S.; Investigation: V.S.D., I.A., A.S., P.R.B., and S.S.; Resources: P.R.B., and S.S.; Writing – original draft: V.S.D., I.A., P.R.B. and S.S.; Writing – reviewing and editing: all authors; Funding acquisition: P.R.B., and S.S.

## Data Availability

All relevant data are included in the paper and/or its Supplementary Information files.

## References

1. Decker, C. J. & Parker, R. P-Bodies and Stress Granules: Possible Roles in the Control of Translation and mRNA Degradation. Cold Spring Harb. Perspect. Biol. 4, a012286–a012286 (2012).

2. Su, X. et al. Phase separation of signaling molecules promotes T cell receptor signal transduction. Science 352, 595–599 (2016).

3. Nozaki, T. et al. Dynamic Organization of Chromatin Domains Revealed by Super-Resolution Live-Cell Imaging. Mol. Cell 67, 282–293.e7 (2017).

4. Pappu, R. V., Cohen, S. R., Dar, F., Farag, M. & Kar, M. Phase Transitions of Associative Biomacromolecules. Chem. Rev. acs.chemrev.2c00814 (2023) doi:10.1021/acs.chemrev.2c00814.

5. Brangwynne, C. P., Tompa, P. & Pappu, R. V. Polymer physics of intracellular phase transitions. Nat. Phys. 11, 899–904 (2015).

6. Brangwynne, C. P. et al. Germline P Granules Are Liquid Droplets That Localize by Controlled Dissolution/Condensation. Science 324, 1729–1732 (2009).

7. Smith, J. et al. Spatial patterning of P granules by RNA-induced phase separation of the intrinsically-disordered protein MEG-3. eLife 5, e21337 (2016).

8. Lasker, K. et al. The material properties of a bacterial-derived biomolecular condensate tune biological function in natural and synthetic systems. Nat. Commun. 13, 5643 (2022).

9. Bowman, G. R. et al. Oligomerization and higher-order assembly contribute to sub-cellular localization of a bacterial scaffold: Mutational analysis of PopZ. Mol. Microbiol. 90, 776–795 (2013).

10. Powers, S. K. et al. Nucleo-cytoplasmic Partitioning of ARF Proteins Controls Auxin Responses in Arabidopsis thaliana. Mol. Cell 76, 177–190.e5 (2019).

11. Davis, R. B., Moosa, M. M. & Banerjee, P. R. Ectopic biomolecular phase transitions: fusion proteins in cancer pathologies. Trends Cell Biol. 32, 681–695 (2022).

12. Kiekebusch, D. & Thanbichler, M. Spatiotemporal organization of microbial cells by protein concentration gradients. Trends Microbiol. 22, 65–73 (2014).

13. Keenan, T. M. & Folch, A. Biomolecular gradients in cell culture systems. Lab Chip 8, 34–57 (2008).

14. Okabe, K. et al. Intracellular temperature mapping with a fluorescent polymeric thermometer and fluorescence lifetime imaging microscopy. Nat. Commun. 3, 705 (2012).

15. Li, Y., Konstantopoulos, K., Zhao, R., Mori, Y. & Sun, S. X. The importance of water and hydraulic pressure in cell dynamics. J. Cell Sci. 133, jcs240341 (2020).

16. Albers, R. W. Biochemical Aspects of Active Transport. Annu. Rev. Biochem. 36, 727–756 (1967).

17. Iacopini, S. & Piazza, R. Thermophoresis in protein solutions. Europhys. Lett. EPL 63, 247–253 (2003).

18. Luetscher, J. A. Biological and Medical Applications of Electrophoresis. Physiol. Rev. 27, 621–642 (1947).

19. Bier, M. Electrophoresis: Theory, Methods, and Applications. (Elsevier Science, Saint Louis, 2013).

20. Anderson, J. L. & Prieve, D. C. Diffusiophoresis: Migration of Colloidal Particles in Gradients of Solute Concentration. Sep. Purif. Methods 13, 67–103 (1984).

21. Prieve, D. C., Anderson, J. L., Ebel, J. P. & Lowell, M. E. Motion of a particle generated by chemical gradients. Part 2. Electrolytes. J. Fluid Mech. 148, 247–269 (1984).

22. Sear, R. P. Diffusiophoresis in Cells: A General Nonequilibrium, Nonmotor Mechanism for the Metabolism-Dependent Transport of Particles in Cells. Phys. Rev. Lett. 122, 128101 (2019).

23. Chong, P. A., Vernon, R. M. & Forman-Kay, J. D. RGG/RG Motif Regions in RNA Binding and Phase Separation. J. Mol. Biol. 430, 4650–4665 (2018).

24. Thandapani, P., O’Connor, T. R., Bailey, T. L. & Richard, S. Defining the RGG/RG Motif. Mol. Cell 50, 613–623 (2013).

25. Alshareedah, I. et al. Interplay between Short-Range Attraction and Long-Range Repulsion Controls Reentrant Liquid Condensation of Ribonucleoprotein–RNA Complexes. J. Am. Chem. Soc. 141, 14593–14602 (2019).

26. Aumiller, W. M. & Keating, C. D. Phosphorylation-mediated RNA/peptide complex coacervation as a model for intracellular liquid organelles. Nat. Chem. 8, 129–137 (2016).

27. Boeynaems, S. et al. Spontaneous driving forces give rise to protein−RNA condensates with coexisting phases and complex material properties. Proc. Natl. Acad. Sci. 116, 7889–7898 (2019).

28. Murthy, A. C. et al. Molecular interactions contributing to FUS SYGQ LC-RGG phase separation and co-partitioning with RNA polymerase II heptads. Nat. Struct. Mol. Biol. 28, 923–935 (2021).

29. Zhou, Q. et al. ATP regulates RNA-driven cold inducible RNA binding protein phase separation. Protein Sci. 30, 1438–1453 (2021).

30. Alshareedah, I., Moosa, M. M., Pham, M., Potoyan, D. A. & Banerjee, P. R. Programmable viscoelasticity in protein-RNA condensates with disordered sticker-spacer polypeptides. Nat. Commun. 12, 6620 (2021).

31. Banerjee, P. R., Milin, A. N., Moosa, M. M., Onuchic, P. L. & Deniz, A. A. Reentrant Phase Transition Drives Dynamic Substructure Formation in Ribonucleoprotein Droplets. Angew. Chem. Int. Ed. 56, 11354–11359 (2017).

32. Alshareedah, I., Moosa, M. M., Raju, M., Potoyan, D. A. & Banerjee, P. R. Phase transition of RNA−protein complexes into ordered hollow condensates. Proc. Natl. Acad. Sci. 117, 15650–15658 (2020).

33. Alshareedah, I., Thurston, G. M. & Banerjee, P. R. Quantifying viscosity and surface tension of multicomponent protein-nucleic acid condensates. Biophys. J. 120, 1161–1169 (2021).

34. Paustian, J. S., Azevedo, R. N., Lundin, S.-T. B., Gilkey, M. J. & Squires, T. M. Microfluidic Microdialysis: Spatiotemporal Control over Solution Microenvironments Using Integrated Hydrogel Membrane Microwindows. Phys. Rev. X 3, 041010 (2013).

35. Shah, P. R. et al. Temperature dependence of diffusiophoresis via a novel microfluidic approach. Lab. Chip 22, 1980–1988 (2022).

36. Jones, D. P. Intracellular diffusion gradients of O_2_ and ATP. Am. J. Physiol.-Cell Physiol. 250, C663– C675 (1986).

37. Shin, S. Diffusiophoretic separation of colloids in microfluidic flows. Phys. Fluids 32, 101302 (2020).

38. Kaur, T. et al. Sequence-encoded and composition-dependent protein-RNA interactions control multiphasic condensate morphologies. Nat. Commun. 12, 872 (2021).

39. Shin, S. et al. Size-dependent control of colloid transport via solute gradients in dead-end channels. Proc. Natl. Acad. Sci. 113, 257–261 (2016).

40. Shin, S., Ault, J. T., Feng, J., Warren, P. B. & Stone, H. A. Low-Cost Zeta Potentiometry Using Solute Gradients. Adv. Mater. 29, 1701516 (2017).

41. Doan, V. S., Chun, S., Feng, J. & Shin, S. Confinement-Dependent Diffusiophoretic Transport of Nanoparticles in Collagen Hydrogels. Nano Lett. 21, 7625–7630 (2021).

42. Palacci, J., Cottin-Bizonne, C., Ybert, C. & Bocquet, L. Osmotic traps for colloids and macromolecules based on logarithmic sensing in salt taxis. Soft Matter 8, 980–994 (2012).

43. Shin, S., Warren, P. B. & Stone, H. A. Cleaning by Surfactant Gradients: Particulate Removal from Porous Materials and the Significance of Rinsing in Laundry Detergency. Phys. Rev. Appl. 9, 034012 (2018).

44. Peter, Q. A. E. et al. Microscale Diffusiophoresis of Proteins. J. Phys. Chem. B 126, 8913–8920 (2022).

45. Annunziata, O., Buzatu, D. & Albright, J. G. Protein Diffusiophoresis and Salt Osmotic Diffusion in Aqueous Solutions. J. Phys. Chem. B 116, 12694–12705 (2012).

46. Doan, V. S., Saingam, P., Yan, T. & Shin, S. A Trace Amount of Surfactants Enables Diffusiophoretic Swimming of Bacteria. ACS Nano 14, 14219–14227 (2020).

47. Shin, S., Doan, V. S. & Feng, J. Osmotic Delivery and Release of Lipid-Encapsulated Molecules via Sequential Solution Exchange. Phys. Rev. Appl. 12, 024014 (2019).

48. Rasmussen, M. K., Pedersen, J. N. & Marie, R. Size and surface charge characterization of nanoparticles with a salt gradient. Nat. Commun. 11, 2337 (2020).

49. Shin, S., Ault, J. T., Warren, P. B. & Stone, H. A. Accumulation of Colloidal Particles in Flow Junctions Induced by Fluid Flow and Diffusiophoresis. Phys. Rev. X 7, 041038 (2017).

50. Palacci, J., Abécassis, B., Cottin-Bizonne, C., Ybert, C. & Bocquet, L. Colloidal Motility and Pattern Formation under Rectified Diffusiophoresis. Phys. Rev. Lett. 104, 138302 (2010).

51. Lafontaine, D. L. J., Riback, J. A., Bascetin, R. & Brangwynne, C. P. The nucleolus as a multiphase liquid condensate. Nat. Rev. Mol. Cell Biol. 22, 165–182 (2021).

52. Riback, J. A. et al. Viscoelastic RNA Entanglement and Advective Flow Underlie Nucleolar Form and Function. http://biorxiv.org/lookup/doi/10.1101/2021.12.31.474660 (2022) xdoi:10.1101/2021.12.31.474660.

53. Abécassis, B., Cottin-Bizonne, C., Ybert, C., Ajdari, A. & Bocquet, L. Boosting migration of large particles by solute contrasts. Nat. Mater. 7, 785–789 (2008).

54. Raynal, F. & Volk, R. Diffusiophoresis, Batchelor scale and effective Péclet numbers. J. Fluid Mech. 876, 818–829 (2019).

55. Florea, D., Musa, S., Huyghe, J. M. R. & Wyss, H. M. Long-range repulsion of colloids driven by ion exchange and diffusiophoresis. Proc. Natl. Acad. Sci. 111, 6554–6559 (2014).

56. Velegol, D., Garg, A., Guha, R., Kar, A. & Kumar, M. Origins of concentration gradients for diffusiophoresis. Soft Matter 12, 4686–4703 (2016).

57. Jung, B., Bharadwaj, R. & Santiago, J. G. Thousandfold signal increase using field-amplified sample stacking for on-chip electrophoresis. Electrophoresis 24, 3476–3483 (2003).

58. Kar, M. et al. Phase-separating RNA-binding proteins form heterogeneous distributions of clusters in subsaturated solutions. Proc. Natl. Acad. Sci. 119, e2202222119 (2022).

59. Fritsch, A. W. et al. Local thermodynamics govern formation and dissolution of Caenorhabditis elegans P granule condensates. Proc. Natl. Acad. Sci. 118, e2102772118 (2021).

60. Chiang, T.-Y. & Velegol, D. Multi-ion diffusiophoresis. J. Colloid Interface Sci. 424, 120–123 (2014).

61. Gupta, A., Rallabandi, B. & Stone, H. A. Diffusiophoretic and diffusioosmotic velocities for mixtures of valence-asymmetric electrolytes. Phys. Rev. Fluids 4, 043702 (2019).

62. Alshareedah, I. et al. Determinants of Viscoelasticity and Flow Activation Energy in Biomolecular Condensates. Sci. Adv. (2024) doi:10.1126/sciadv.adi6539.

63. Young, N. O., Goldstein, J. S. & Block, M. J. The motion of bubbles in a vertical temperature gradient. J. Fluid Mech. 6, 350–356 (1959).

64. Guo, W. et al. Non-associative phase separation in an evaporating droplet as a model for prebiotic compartmentalization. Nat. Commun. 12, 3194 (2021).

65. Ianeselli, A. et al. Non-equilibrium conditions inside rock pores drive fission, maintenance and selection of coacervate protocells. Nat. Chem. 14, 32–39 (2022).

66. Testa, A. et al. Sustained enzymatic activity and flow in crowded protein droplets. Nat. Commun. 12, 6293 (2021).

67. Demarchi, L., Goychuk, A., Maryshev, I. & Frey, E. Enzyme-Enriched Condensates Show Self-Propulsion, Positioning, and Coexistence. Phys. Rev. Lett. 130, 128401 (2023).

68. Agrawal, A., Ganai, N., Sengupta, S. & Menon, G. I. Nonequilibrium Biophysical Processes Influence the Large-Scale Architecture of the Cell Nucleus. Biophys. J. 118, 2229–2244 (2020).

69. Ramm, B. et al. A diffusiophoretic mechanism for ATP-driven transport without motor proteins. Nat. Phys. 17, 850–858 (2021).

70. Shim, S. et al. CO_2_-Driven diffusiophoresis for maintaining a bacteria-free surface. Soft Matter 17, 2568–2576 (2021).

71. Vrhovec Hartman, S., Božič, B. & Derganc, J. Migration of blood cells and phospholipid vesicles induced by concentration gradients in microcavities. New Biotechnol. 47, 60–66 (2018).

72. Hajian, R. & Hardt, S. Formation and lateral migration of nanodroplets via solvent shifting in a microfluidic device. Microfluid. Nanofluidics 19, 1281–1296 (2015).

73. Deamer, D. Concentration Gradients. in Encyclopedia of Astrobiology (eds. Gargaud, M.et al.) 354–355 (Springer Berlin Heidelberg, Berlin, Heidelberg, 2011). doi:10.1007/978-3-642-11274-4_226.

