## Supplementary Information for "Diffusiophoresis promotes phase separation and transport of biomolecular condensates"

### MATERIALS AND METHODS

#### Microfluidic device fabrication

The channel was created using the microfluidic sticker technique with UV-curable epoxy (Norland Optical Adhesive, NOA-81).[1] In this method, a polydimethylsiloxane (PDMS, Dow Inc.) sticker is cast from the SU-8 mold. Subsequently, the NOA-81 channel is molded from the PDMS sticker and partially cured under UV light (IntelliRay 400, Uvitron). Another flat piece of NOA-81 is then combined with the molded channel and exposed to UV light for complete curing. Once the channel is formed, a solution of polyethylene glycol diacrylate (PEGDA, Sigma-Aldrich) (20%(v/v) PEGDA, 2%(v/v) photoinitiator 2-hydroxy-2-methylpropiophenone, Sigma-Aldrich) is injected into the empty epoxy channel. The PEGDA membrane is selectively cured within a thin region. To generate a thin band of UV light, a photomask containing a thin transparent strip and a 40× objective lens is used to focus the parallel light.[2–4] After the thin membrane is cured near the inlet, any uncured PEGDA is flushed out by deionized (DI) water for 30 minutes (**Fig. S1**).

#### Sample preparation

ssDNA oligos of length 40 were purchased from Integrated DNA Technologies (NJ, USA). The dry stocks were reconstituted in RNase-free water. The reconstituted solutions were centrifuged at 23,000 × g for 2 minutes to remove any particles. After extracting the supernatant, DNA concentrations were subsequently measured using a NanoDrop 1C™ spectrophotometer. The DNA stocks were then aliquoted and stored at −20°C for further use. The peptide $[RGRGG]_5$ was synthesized by Genscript Inc, USA., and was reconstituted in RNase-free water containing 50 mM DTT (dithiothreitol, ThermoFisher Scientific). The purity of the peptides was higher than 90% as per manufacturer specifications. All the peptide sequences contain a C-terminal cysteine that is used for site-specific labeling with fluorescence dyes. The site-specific peptide labeling was performed as described previously.[5,6] Polyuridylic acid [poly(rU); molecular weight = 600-1000 kDa] was purchased from Sigma-Aldrich. Poly(rU) RNA was re-suspended in RNase-free water at concentrations ≥ 80 mg/ml, then aliquots were made and stored at −20°C for later use. All the ssDNA, RNA, and peptide stocks were checked for complete solubilization and absence of aggregates under the microscope. Salmon protamine (P4005) was purchased from Sigma-Aldrich and used without any further purification.

#### Microscopy visualization of diffusiophoresis and phase separation

The entire channel was first flushed with the solution consisting of salt, protein, Tris-HCl (10 mM), and DTT (20 mM) in RNase-free water using a syringe pump (Pump 11 Pico Plus Elite, Harvard Apparatus). Subsequently, a second solution consisting of DNAs at different salt concentrations was injected into the left side channel with a flow rate of 20 μl/hr to initiate the phase separation experiment in the salt gradient. The transport and phase separation dynamics were observed under an inverted fluorescence microscope (DMi8, Leica) equipped with an sCMOS camera (ORCA-Flash4.0 LT3, Hamamatsu). The recorded images were analyzed using ImageJ and MATLAB. For the visualization of the condensates, Alexa488-labeled peptide $[RGRGG]_5$ and Cy5-labeled ssDNA $[dT]_{40}$ (purchased from Integrated DNA Technologies (NJ, USA)) were used.

#### Estimation of condensate size distribution

To determine the condensate size distribution using confocal microscopy, the condensates at three different salt concentrations were prepared by mixing 0.5 mg/ml peptide $[RGRGG]_5$ and 0.625 mg/ml ssDNA $[dT]_{40}$ in a buffer containing 10 mM Tris-HCl (pH 7.5), 20 mM DTT, and either 1, 10, or 20 mM

NaCl with 150 nM each of Alexa488-labeled $[RGRGG]_5$ and Cy5-labeled $[dT]_{40}$. 5.0 μl of the condensate forming samples were drop-casted on a Tween20 [20%(v/v)] coated coverslips (0.17 mm thickness, 18x18 $mm^2$ dimensions). The sample was then sandwiched using a glass slide, creating a sample chamber using double-sided tape strips. 100 μl of mineral oil was injected into the sample chamber to prevent evaporation by filling the vacant space in the sample chamber. A laser scanning confocal microscope (Q2, ISS Inc.) with a 60x water objective was used for acquiring the fluorescence images of the condensates. The condensates in the fluorescence images were approximated as circles using Hough Circle Transform algorithm under UCB Vision Sciences plugin in ImageJ (https://imagej.net/plugins/hough-circle-transform).[7] The diameters of the circles were determined, and a distribution of the estimated diameters was plotted for 1 mM, 10 mM, and 20 mM NaCl concentration (**Figs. S2a,b**).

The size distribution was also measured using dynamic light scattering (Lite Sizer 500, Anton Paar). The condensates were formed with the same mixing conditions as the above confocal microscopy measurement. The sample with a total volume of 60 μl was loaded into a low-volume cuvette (Univette). The measurement was initiated immediately after forming the condensates to prevent sedimentation. The sizes of the condensates were obtained using the General analysis mode and Advanced cumulant model. The size distribution is reported in Fig. S2c with different NaCl concentrations, ranging from 1 mM to 20 mM.

**Zeta potential measurements of biomolecular condensates**

Zeta potential measurements (Lite Sizer 500, Anton Paar) of two types of biomolecular condensates ($[dT]_{40}$- $[RGRGG]_5$ and poly(rU)-protamine) are presented in **Fig. S3**. Samples were prepared by mixing the protein ($[RGRGG]_5$ or protamine) and the nucleic acid ($[dT]_{40}$ ssDNA or poly(rU) RNA) at the desired concentration ratio in a buffer containing 10 mM Tris-HCl. For $[dT]_{40}$-$[RGRGG]_5$ mixtures, the peptide concentration was fixed at 0.5 mg/ml while the DNA concentration was varied. For poly(rU)-protamine mixtures, the protein concentration was fixed at 0.15 mg/ml while the RNA concentration was varied. Next, the sample was placed in a cuvette and inserted into Litesizer 500 (Anton Paar) particle size analyzer to perform dynamic light scattering measurements. We observe a charge inversion of protein-nucleic acid complex upon increasing the mixing ratio. $[dT]_{40}$-rich droplets (ssDNA-to-protein ratio $n_D/n_P$ = 1.25) have a negative charge (zeta potential = –27.2 ± 1.5 mV), while $[RGRGG]_5$-rich biomolecular condensates show a weakly positive charge (zeta potential = 9.8 ± 1 mV at $n_D/n_P$ = 1/9). On the other hand, poly(rU)-rich condensates (RNA-to-protein ratio $n_R/n_P$ = 4) have a zeta potential of –41.7 ± 1 mV while the charge of the condensates is reversed (zeta potential = 58.8 ± 1 mV) as the concentration shifts to the protamine-rich region ($n_R/n_P$ = 1/4).

**Salt-dependent diffusiophoretic mobilities of biomolecules**

While diffusiophoretic mobility is often treated as constant,[8] it is a property that is sensitive to the local zeta potential, thus the local salt concentration. Due to the small size of the biomolecules, the contribution of chemiphoresis can be effectively neglected. Therefore, diffusiophoretic mobility $M_i$ is directly proportional to the zeta potential of the protein

$$M_i = \zeta \frac{\varepsilon}{\eta} \frac{k_B T}{ze} \beta \tag{S1}$$

where $\varepsilon$ is the permittivity, $\eta$ is the viscosity, $k_B$ is the Boltzmann constant, $T$ is the temperature, $z$ is the valence, and $e$ is the element charge and $\beta = (D_+ - D_-)/(D_+ + D_-)$, is the diffusivity contrast, where $D_+$

and $D_-$ are the diffusivity of cations and anions, respectively.[9] For weakly charged particles, the zeta potential scales with the local salt concentration $c$ as $\zeta \sim 1/\sqrt{c}$.[10] Hence, equation S1 can be rewritten as

$$M_i = \zeta_0 \sqrt{\frac{c_0}{c}} \frac{\varepsilon}{\eta} \frac{k_B T}{Ze} \beta \quad \text{(S2)}$$

where $\zeta_0$ is the zeta potential at $c_0$ = 1 mM. The concentration of solutes within the channel depends both on position and time $c = c(x,t)$, which can be determined by solving the one-dimensional diffusion equation $\partial_t c = D_s \partial_{xx} c$, where $D_s = 2D_+ D_- /(D_+ + D_-)$ is the ambipolar diffusion coefficient. Here, instead of accounting for the full solute concentration distribution for determining the concentration-dependent $M_i$, we make an approximation where we average the local concentration across the channel at a given time such that the averaged concentration only depends on time, shown as

$$c = \bar{c}(t) = \frac{1}{L} \int_{x=0}^{L} c(x,t) dx. \quad \text{(S3)}$$

This allows us to approximate the concentration-dependent mobility (which is position and time-dependent) as only time-dependent

$$M_i(t) = \zeta_0 \sqrt{\frac{c_0}{\bar{c}(t)}} \frac{\varepsilon}{\eta} \frac{k_B T}{Ze} \beta. \quad \text{(S4)}$$

This position-averaged mobility is used to plot **Figs. 3b,d** in the main text and **Fig. S5**. The time-averaged mobility values $\langle M \rangle$ reported in **Fig. 3e** are the mobility in equation S4 time-averaged over the course of the experiments.

**Turbidity measurements**

Samples were prepared in a tube by mixing the peptide and the ssDNA at the desired mixing ratio and a fixed peptide concentration of 1.0 mg/ml. The buffer of these samples contains 10 mM Tris-HCl (pH 7.5) and 20 mM NaCl. The sample was then placed on a UV-Vis spectrophotometer (NanoDrop 1C) and the solution turbidity at 350 nm was measured for three independent samples. Before measuring the turbidity of peptide-ssDNA samples, the instrument was blanked using the experimental buffer. The values of the turbidity were averaged for each mixing ratio, and the error was estimated as half the range of experimentally measured values (**Fig. S6**).

**Diffusiophoresis in $[RGRGG]_5$-rich and poly(rU)-rich condensates**

We have investigated the salt gradient effects in other biomolecular systems, e.g., different stoichiometry or species. For instance, we studied $[RGRGG]_5$-rich $[dT]_{40}$-$[RGRGG]_5$ condensates by introducing $[RGRGG]_5$ at an elevated concentration of 0.9 mg/ml in the left reservoir while lowering the concentration of $[dT]_{40}$ to 0.15 mg/ml in the center and right channels (**Figs. S7a,b**). Because of the opposite configuration, we choose potassium acetate (KAc) as the solute, which can drive $[RGRGG]_5$ up the gradient and $[dT]_{40}$ down the gradient due to the positive $\beta$ factor ($\beta$ = 0.29) such that diffusiophoresis is expected to accelerate their migration toward each other. We observe that the KAc gradients ($c_1$ = 20 mM, $c_2$ = 1 mM) resulted in a stronger condensate formation, but does not significantly affect the transport of the formed condensates (**Figs. S7a,b**). This is because KAc gradients can accelerate the transport of individual $[RGRGG]_5$ and $[dT]_{40}$ molecules (**Fig. S4**), a stronger condensate formation is expected. However, the condensates do not experience any significant active motility due to the low zeta potential ($\zeta$ = 9.5 mV; **Fig. S3**) of $[RGRGG]_5$-rich condensates. We note that $[dT]_{40}$-$[RGRGG]_5$ mixtures display an asymmetric charge inversion behavior (**Fig. S3**) where the $[dT]_{40}$-rich condensates carry more charge than the $[RGRGG]_5$-rich condensates, similar to previously reported poly(rU)-$[RP]_3$ system,[11] and poly(rA)-$[RGRGG]_5$ condensates.[5]

The poly(rU)-protamine condensates were formed by introducing poly(rU) in the left reservoir (1.2 mg/ml) and protamine initially in the center and right channels (0.3 mg/ml). Using NaCl, we observe similar behavior to $[dT]_{40}$-rich $[dT]_{40}$-$[RGRGG]_5$ condensates, where the negatively charged condensates were formed and the transport of poly(rU)-protamine condensates was improved under NaCl gradients ($c_1$ = 20 mM and $c_2$ = 1 mM) (**Fig. S7c**) in comparison with the case without NaCl gradient ($c_1$ = $c_2$ = 20 mM; **Fig. S7d**). This is due to the high negative surface charge ($\zeta$ = –61.5 mV at RNA-to-protein ratio 4; **Fig. S3**) of the poly(rU)-rich condensates.

#### SUPPLEMENTARY FIGURES

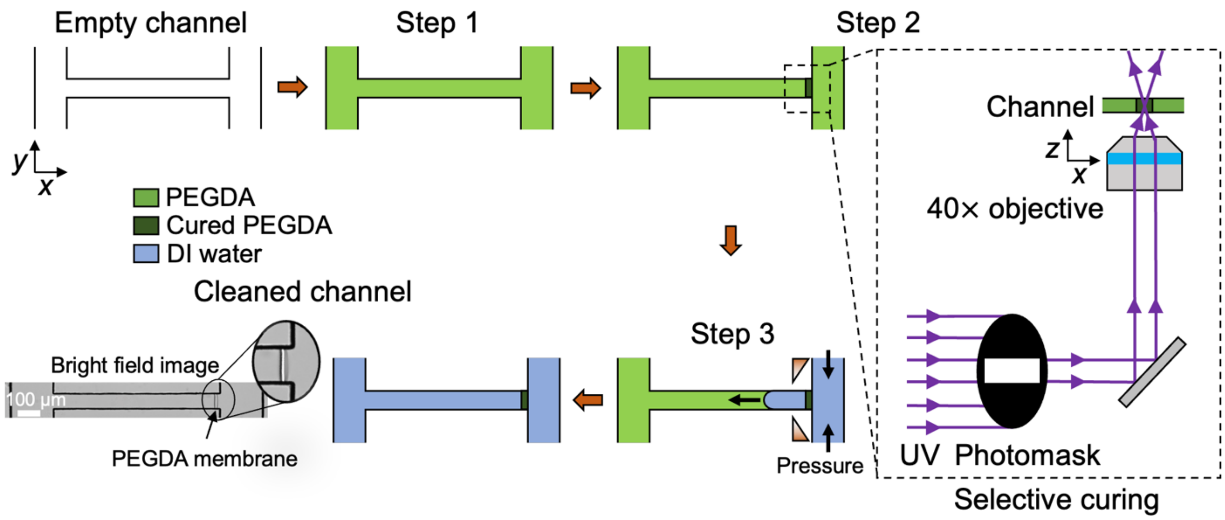


**Figure S1.** In situ fabrication of PEGDA membranes. In Step 1, the empty channel was filled with PEGDA solution [20% (v/v) PEGDA, 2% (v/v) photoinitiator 2-hydroxy-2-methylpropiophenone]. Then, a black mask with a thin transparent strip and 40× objective were used to selectively cure a thin region to create a PEGDA membrane (Step 2). Finally, the excessive PEGDA was flushed out through the membrane by DI water at elevated pressure (Step 3). The bright field image shows the PEGDA membrane near the right inlet of the channel.

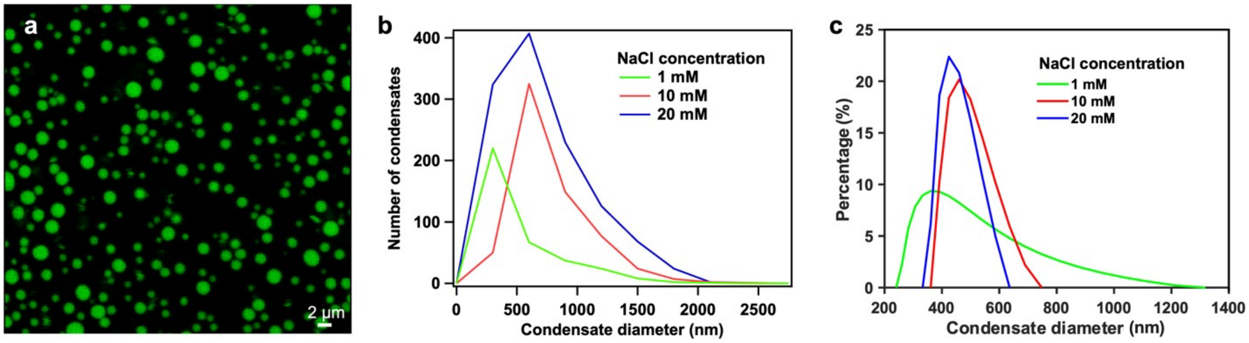


**Figure S2.** Size distribution of $[dT]_{40}$-$[RGRGG]_5$ condensates ($[dT]_{40}$:$[RGRGG]_5$ = 1.25:1). (**a**) Confocal image of $[dT]_{40}$-$[RGRGG]_5$ condensates in 20 mM NaCl (sample composition: 0.5 mg/ml $[RGRGG]_5$, 150 nM Alexa-488 labeled $[RGRGG]_5$, 0.625 mg/ml $[dT]_{40}$, 10 mM Tris-HCl, 20 mM DTT, and 20 mM NaCl). (**b**, **c**) Size distribution of $[dT]_{40}$-$[RGRGG]_5$ condensates at varying NaCl concentrations obtained from the (**b**) confocal imaging and (**c**) dynamic light scattering.

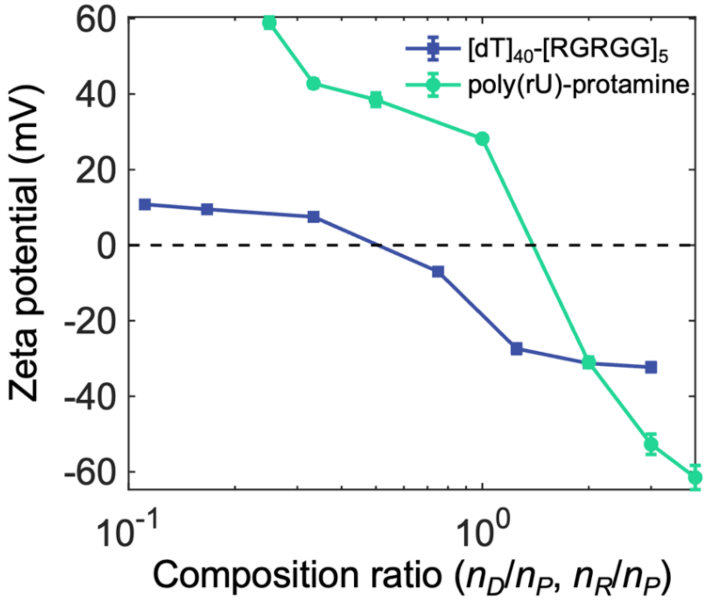


**Figure S3.** Zeta potential of $[dT]_{40}$-$[RGRGG]_5$ and poly(rU)-protamine complexes with varying composition ratios (ssDNA/RNA-to-protein; $n_{\{D,R\}}/n_P$). Error bars represent standard deviation.

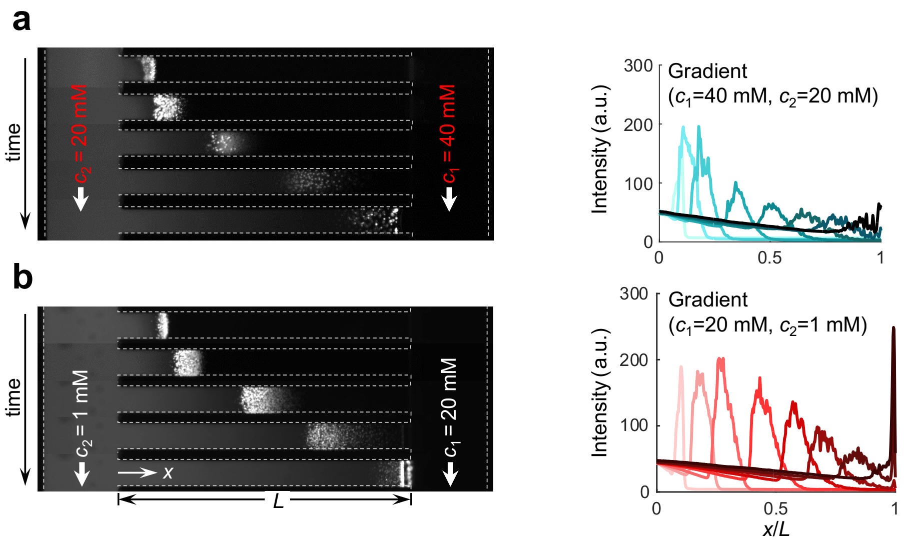


**Figure S4.** Formation and transport of the condensates under different salt contrasts. The image sequences and their corresponding width-averaged intensity distribution of the formation and transport $[dT]_{40}$-$[RGRGG]_5$ condensates over 60 minutes. The NaCl concentrations are (**a**) $c_1$ = 40 mM, $c_2$ = 20 mM, and (**b**) $c_1$ = 20 mM, $c_2$ = 1 mM. The concentration difference is nearly identical in both cases (~20 mM). However, the formation and transport of the condensates in (**b**) are stronger than in (**a**) due to the logarithmic dependence of diffusiophoresis.

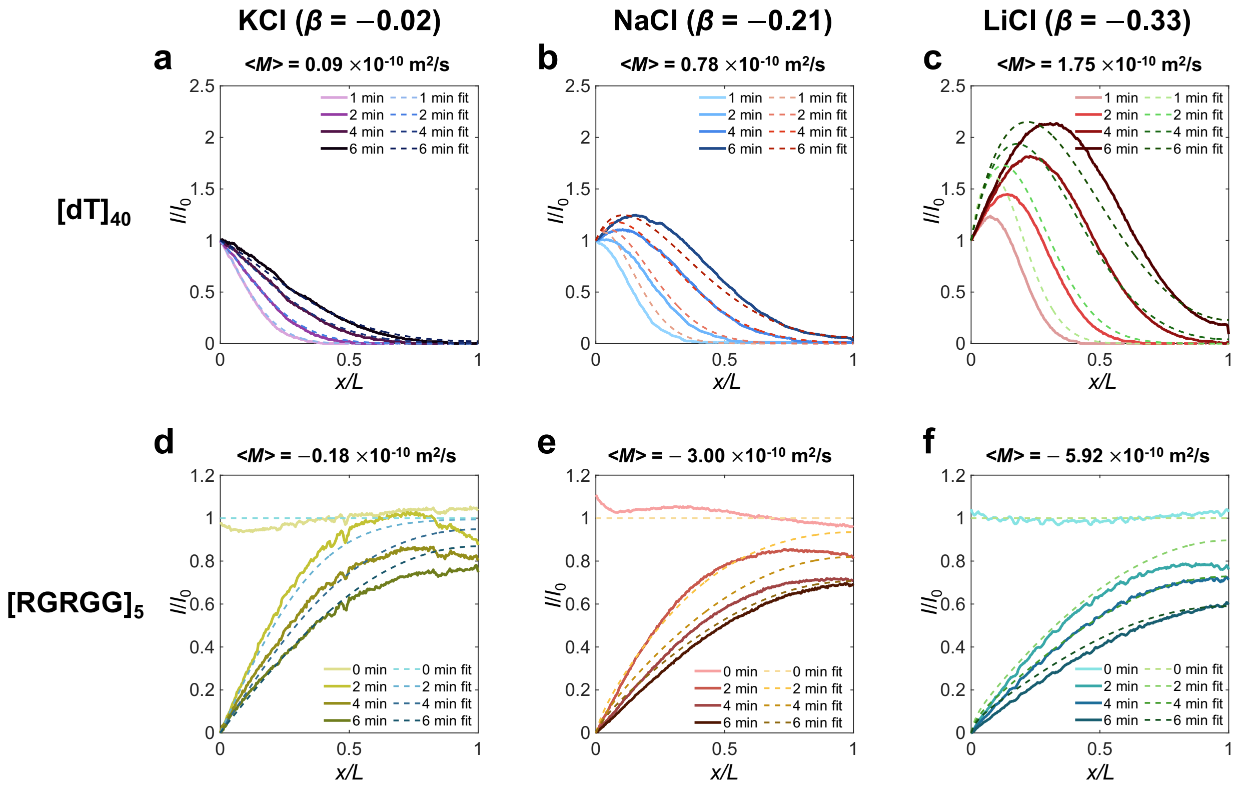


**Figure S5.** The diffusiophoretic mobility of $[dT]_{40}$ and $[RGRGG]_5$ in various types of salts (varying $\beta$). (**a-c**) The mobility of $[dT]_{40}$ and (**d-f**) $[RGRGG]_5$ in KCl, NaCl and LiCl gradient, respectively. The highest mobilities were observed in LiCl (**c**,**f**), while the values were the lowest in the KCl gradient (**a**,**d**). These differences in mobilities across different salts suggest that the transport of $[dT]_{40}$ and $[RGRGG]_5$ is influenced by diffusiophoresis, which is dependent on the diffusivity contrast between ions resulting from the varying salt gradients.

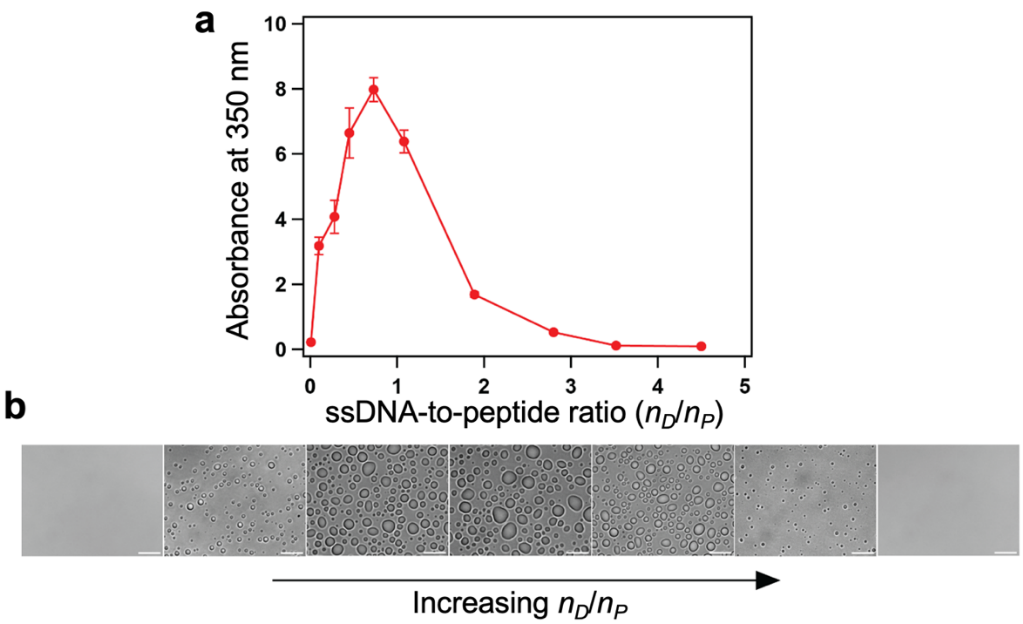


**Figure S6.** (**a**) A plot showing absorbance at 350 nm (turbidity) of $[dT]_{40}$-$[RGRGG]_5$ mixtures at different mixing stoichiometries. Error bars represent standard deviation. (**b**) Brightfield images of the condensates formed at increasing ssDNA-peptide selected mixing ratios for which turbidity measurements were performed in (**a**). The scale bar is 10 μm. From both (**a**) and (**b**) reentrant phase separation of $[dT]_{40}$-$[RGRGG]_5$ condensates is evident.

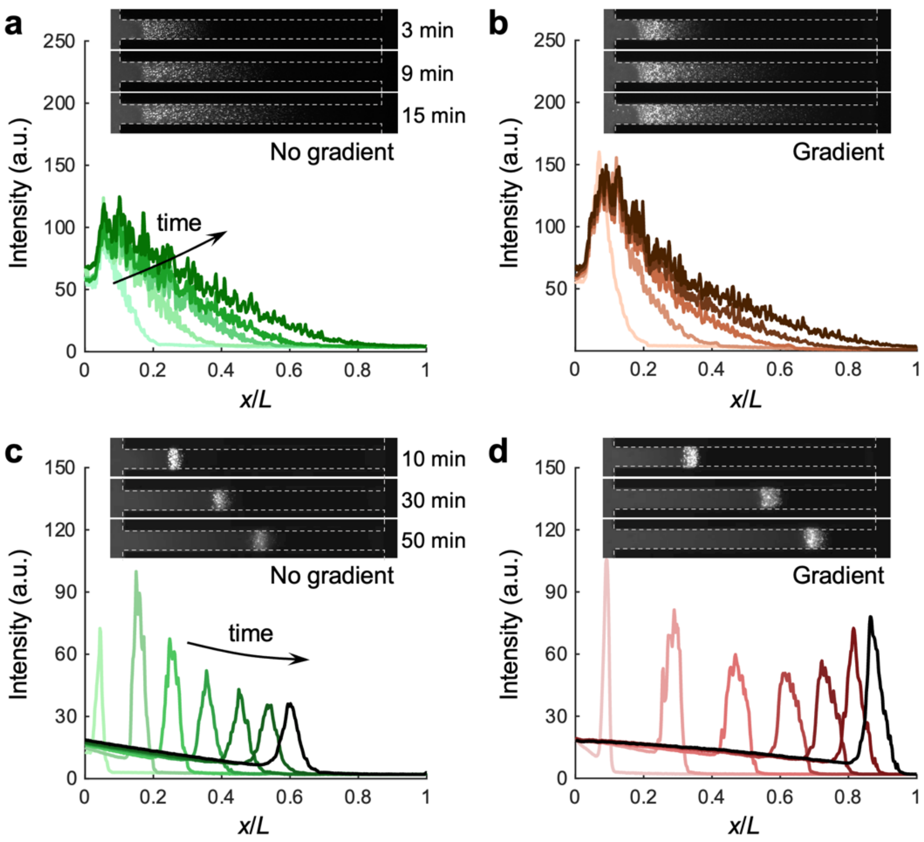


**Figure S7.** Fluorescence images of (**a,b**) [RGRGG]$_5$-rich [dT]$_{40}$-[RGRGG]$_5$ condensates (**a**) without ($c_1$ = $c_2$ = 20 mM) or (**b**) with KAc gradients ($c_1$ = 20 mM and $c_2$ = 1 mM), and (**c,d**) poly(rU)-rich poly(rU)-protamine condensates (**c**) without ($c_1$ = $c_2$ = 20 mM) or (**b**) with NaCl gradients ($c_1$ = 20 mM and $c_2$ = 1 mM). The timestamps for the intensity profiles for (**a,b**) are 1, 3, 6, 9, and 15 min, and (**c,d**) are 1, 10, 20, 30, 40, 50, and 60 min, respectively.

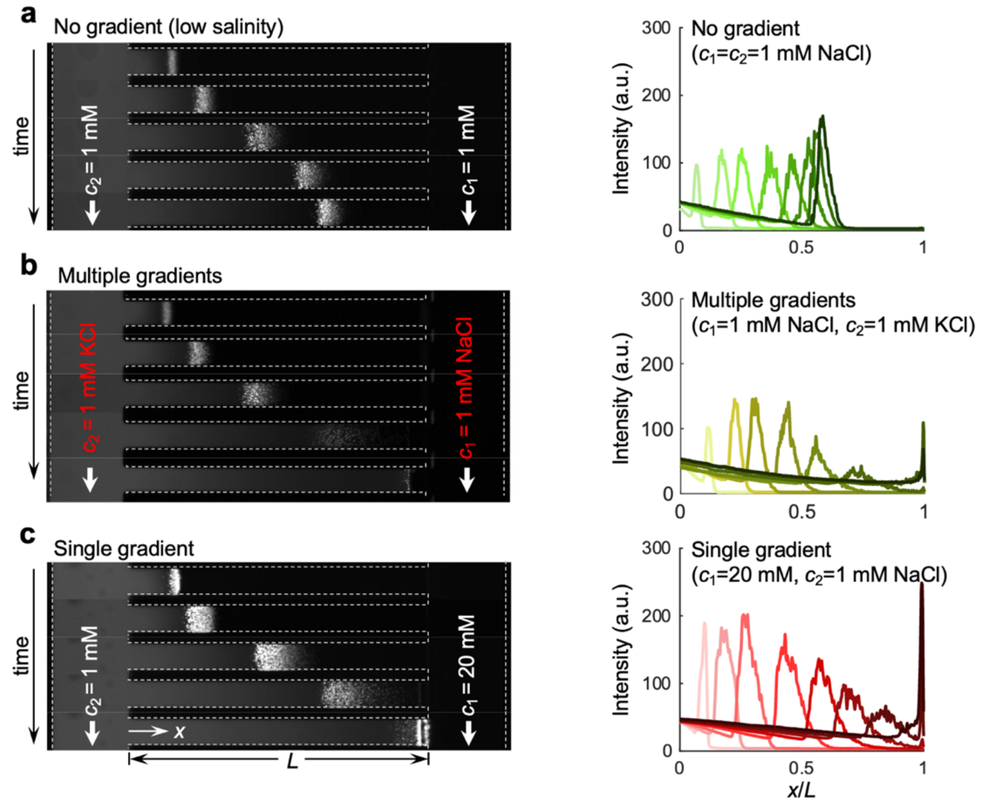


**Figure S8.** Formation and transport of condensates by interdiffusion of equimolar multispecies solutes. (**a**) The image sequences the formation and transport $[dT]_{40}$-$[RGRGG]_5$ condensates over a 60-minute period (**a**) without any salt gradient ($c_1 = c_2$ = 10 mM NaCl); (**b**) under interdiffusion of equimolar NaCl and KCl ($c_1$ = 1 mM NaCl, $c_2$ = 1 mM KCl) and (**c**) under NaCl gradient ($c_1$ = 20 mM, $c_2$ = 1 mM). In the case of interdiffusistaton of equimolar NaCl and KCl in (**b**), due to the difference in the diffusion coefficients of $Na^+$ and $K^+$, a local electrical field arises, causing electrophoresis. As a result, the transport and the formation of the condensate are improved compared to the case of no salt gradients (**a**). However, the lack of chemiphoresis in (**b**) due to the equimolar concentration leads to relatively weaker improvement in the transport and formation compared to the single NaCl gradient case in (**c**).

#### SUPPLEMENTARY MOVIE LEGENDS

**Movie S1.** Formation and transport of $[dT]_{40}$ and $[RGRGG]_5$ condensates visualized using Cy5-labeled $[dT]_{40}$ in the presence and absence of the NaCl gradients. The condensates were observed under a 10× objective. Top: $c_1 = c_2 = 20$ mM; middle: $c_1 = c_2 = 1$ mM; bottom: $c_1 = 20$ mM, $c_2 = 1$ mM.

**Movie S2.** Phase separation, transport, and dissolution of $[dT]_{40}$-$[RGRGG]_5$ condensates observed under a 40× objective. Top: $c_1 = c_2 = 20$ mM; middle: $c_1 = c_2 = 1$ mM; bottom: $c_1 = 20$ mM, $c_2 = 1$ mM.

#### SUPPLEMENTARY REFERENCES


1. Bartolo, D., Degré, G., Nghe, P. & Studer, V. Microfluidic stickers. *Lab Chip* **8**, 274–279 (2008).
2. Shah, P. R. *et al.* Temperature dependence of diffusiophoresis *via* a novel microfluidic approach. *Lab Chip* **22**, 1980–1988 (2022).
3. Nery-Azevedo, R., Banerjee, A. & Squires, T. M. Diffusiophoresis in Ionic Surfactant Gradients. *Langmuir* **33**, 9694–9702 (2017).
4. Paustian, J. S., Azevedo, R. N., Lundin, S.-T. B., Gilkey, M. J. & Squires, T. M. Microfluidic Microdialysis: Spatiotemporal Control over Solution Microenvironments Using Integrated Hydrogel Membrane Microwindows. *Phys. Rev. X* **3**, 041010 (2013).
5. Alshareedah, I. *et al.* Interplay between Short-Range Attraction and Long-Range Repulsion Controls Reentrant Liquid Condensation of Ribonucleoprotein–RNA Complexes. *J. Am. Chem. Soc.* **141**, 14593–14602 (2019).
6. Kaur, T. *et al.* Sequence-encoded and composition-dependent protein-RNA interactions control multiphasic condensate morphologies. *Nat. Commun.* **12**, 872 (2021).
7. D'Orazio, T., Guaragnella, C., Leo, M. & Distante, A. A new algorithm for ball recognition using circle Hough transform and neural classifier. *Pattern Recognit.* **37**, 393–408 (2004).
8. Shin, S. *et al.* Size-dependent control of colloid transport via solute gradients in dead-end channels. *Proc. Natl. Acad. Sci.* **113**, 257–261 (2016).
9. Peter, Q. A. E. *et al.* Microscale Diffusiophoresis of Proteins. *J. Phys. Chem. B* **126**, 8913–8920 (2022).
10. Kirby, B. J. & Hasselbrink, E. F. Zeta potential of microfluidic substrates: 1. Theory, experimental techniques, and effects on separations. *Electrophoresis* **25**, 187–202 (2004).
11. Banerjee, P. R., Milin, A. N., Moosa, M. M., Onuchic, P. L. & Deniz, A. A. Reentrant Phase Transition Drives Dynamic Substructure Formation in Ribonucleoprotein Droplets. *Angew. Chem. Int. Ed.* **56**, 11354–11359 (2017).
